# Central inhibition of P2Y_12_R differentially regulates survival and neuronal loss in MPTP-induced Parkinsonism in mice

**DOI:** 10.1101/2021.02.03.256941

**Authors:** András Iring, Adrián Tóth, Mária Baranyi, Lilla Otrokocsi, László V. Módis, Flóra Gölöncsér, Bernadett Varga, Tibor Hortobágyi, Dániel Bereczki, Ádám Dénes, Beáta Sperlágh

**Author notes:** Correspondence should be sent to: Beáta Sperlágh, Laboratory of Molecular Pharmacology, Institute of Experimental Medicine, 1083 Budapest, Hungary.

## Abstract

Parkinson’s disease (PD) is a chronic, progressive neurodegenerative condition; characterized with the degeneration of the nigrostriatal dopaminergic pathway and neuroinflammation. During PD progression, microglia, the resident immune cells in the central nervous system (CNS) display altered activity, but their role in maintaining PD development has remained unclear to date. The purinergic P2Y12 receptor (P2Y12R), which is exclusively expressed on the microglia in the CNS has been shown to regulate microglial activity and responses; however, the function of the P2Y12R in PD is unknown. Here we show that while pharmacological or genetic targeting of P2Y12R previous to disease onset augments acute mortality, these interventions protect against neurodegenerative cell loss and the development of neuroinflammation in vivo. Pharmacological inhibition of receptors during disease development reverses the symptoms of PD and halts disease progression. We found that P2Y12R regulate ROCK and p38 MAPK activity and control cytokine production. Understanding protective and detrimental P2Y12 receptor-mediated actions in the CNS may reveal novel approaches to control neuroinflammation and modify disease progression in PD.

## Introduction

Parkinson’s disease (PD) is a chronic, progressive neurodegenerative disease, initially described as early as in 1817 (1). The incidence of new Parkinson’s cases ranges from 5 to 15 in 100,000 annually (2, 3), most prominently during the sixth to ninth decades of life. PD patients suffer debilitating symptoms that reduce the quality of life and present increased mortality risks (4, 5); symptoms that include motor function impairments, such as asymmetric tremor, cogwheel rigidity, bradykinesia, as well as non-motor features, including constipation, depression and sleep disorder (2, 6). Characteristic pathological feature of PD is the selective degeneration of dopaminergic neurons in the substantia nigra *pars compacta* and deposition of Lewy-bodies, which are cytoplasmic aggregates of insoluble α-synuclein. While both genetic and environmental factors influence disease development, they converge on common pathways to induce PD. These pathways include mitochondrial dysfunction, oxidative stress, impaired autophagy, protein aggregation and neuroinflammation (7). Inflammatory processes developing in the central nervous system (CNS) contribute to brain development as well as neuropathological events (8, 9). Neurodegeneration during PD occurs either through “cell-autonomous” mechanisms or via indirect degeneration caused by interaction with resident glial cells (astrocytes, microglia). Microglia are the primary innate immune cells in the CNS, continually monitoring synaptic activity, clearing apoptotic cells, and reacting to pathological events through complex inflammatory responses (8). Microglial activation has been implicated in the pathomechanism of PD in humans (10); however, it is unclear whether microglia are involved in the initiation of dopaminergic cell death in PD or increased microglial activity merely responds to the production of cytokines or damage-associated molecules from degraded neurons. Nonetheless, several pro-inflammatory cytokines were found elevated in the substantia nigra of postmortem human samples, such as IL-1β, IL-2, IL-6 and tumor necrosis factor α (TNFα) (11, 12), as well as anti-inflammatory mediators, namely transforming growth factor β1 or IL-10 (13). Several studies demonstrate functional differences between monocyte-derived macrophages and microglia regarding immunoregulatory, cell migratory and phagocytic function (14–16). These observations highlight the distinct properties of the myeloid lineage and imply that microglia contribute differently from other myeloid cells to CNS injury and repair. One particular feature of microglia is that the G-protein coupled purinergic receptor P2Y_12_ (P2Y_12_R) is expressed on their cell body and ramified processes, allowing the discrimination of microglia from other cells of the myeloid lineage (17).

The P2Y_12_R is activated by adenosine diphosphate (ADP), and activation of the receptor promotes the dissociation of the Gα_i_ subunit, which in turn reduces adenylate-cyclase activity and intracellular cAMP levels. Our previous work implicated P2Y_12_R in the regulation of inflammatory pain, where receptor inhibition decreased pro-inflammatory IL-1β, TNFα and CXCL1 levels (18). Recently, we have identified novel microglia-neuron somatic interaction sites and the regulatory role of P2Y_12_R in neuronal activation and neuroprotection was also demonstrated (19). Microglia isolated from P2Y_12_R-deficient mice show normal baseline motility, but have impaired polarization, migration or branch extension towards sites of injury (20). While the involvement of P2Y_12_R has been explored in several pathological conditions, the function of the receptor in Parkinson’s disease has not been studied yet.

Presently available treatment options for Parkinson’s disease are based on the supplementation of levodopa (3,4-dihydroxy-l-phenylalanine), an intermediate in the dopamine synthesis pathway, and augmented with drugs that enhance the efficiency of levodopa, such as aromatic L-amino acid decarboxylase inhibitors, catechol-O-methyltransferase blockers and monoamine oxidase type B inhibitors (21). The shortcoming of these therapies is that the treatment is symptomatic, improving mainly the motor impairments, moreover, has no effect on disease progression.

Here we show that the neuroinflammatory aspect of the disease is closely mimicked by MPTP administration in mice. We have observed that P2Y_12_-receptors are necessary for the activation of microglia and are involved in the progression of the disease. Interestingly, blockade of the P2Y_12_-receptor augments acute mortality after MPTP administration, but protects against the neurodegenerative cell loss and hinders neuroinflammation during experimental PD. Additionally, pharmacological inhibition of the receptor during disease progression alleviates the symptoms of Parkinson’s disease by modulating Rho-kinase (ROCK) and p38 MAPK activity and reducing pro- and anti-inflammatory cytokine production. Interestingly, while ROCK-inhibition via fasudil has an apparent protective effect during experimental PD, it also conveys a wide-range of undesirable side-effects that are detrimental during disease resolution. Our results reveal the involvement of centrally expressed P2Y_12_R during initiation and progression of experimentally-induced Parkinson’s disease, and propose the receptor as a promising highly selective and specific pharmacological target to impede disease progression in PD.

## Results

### Constitutive genetic deletion of the P2Y_12_ receptor promotes mortality, but preserves dopaminergic neurons and alleviates PD-like symptoms after MPTP-administration

First, we investigated the role of P2Y_12_ receptors (P2Y_12_R) in the development of Parkinson’s disease, utilizing mice with genetic deficiency for the *P2yr12* gene (herein referred to as P2Y_12_R-KO) or using pharmacological blockade by PSB 0739, a specific P2Y_12_R inhibitor. We have observed that genetic deletion or pharmacological blockade of the receptor contributes to lower survival following MPTP-treatment (Figure 1A). While the survival rate in wild-type control and P2Y_12_R-KO mice receiving only PSB 0739 was indistinguishable from the vehicle treated control group (data not shown), MPTP administration contributed to a casualty rate of approximately 25 % in wild-type control mice. Remarkably, MPTP treatment combined with dysfunctional P2Y_12_R markedly increased mortality (approximately 41 % in wild-type PSB 0739 group (*P=0.0242 vs. WT control*) and 45 % in P2Y_12_R-KO groups (*P=0.0052 vs. WT control)*). This observation emphasizes that the presence of functional P2Y_12_R is necessary to prevent the deterioration of acute neurotoxic events and to initiate disease resolution after MPTP treatment; similarly to the reported role of P2Y_12_R expressed on microglia after viral infection or ischaemic brain injury (22, 23). Microglia-depletion has also been shown to exacerbate Parkinson’s disease symptoms and augments the infiltration of leukocytes into the brain after MPTP treatment (24).

**Figure 1.**
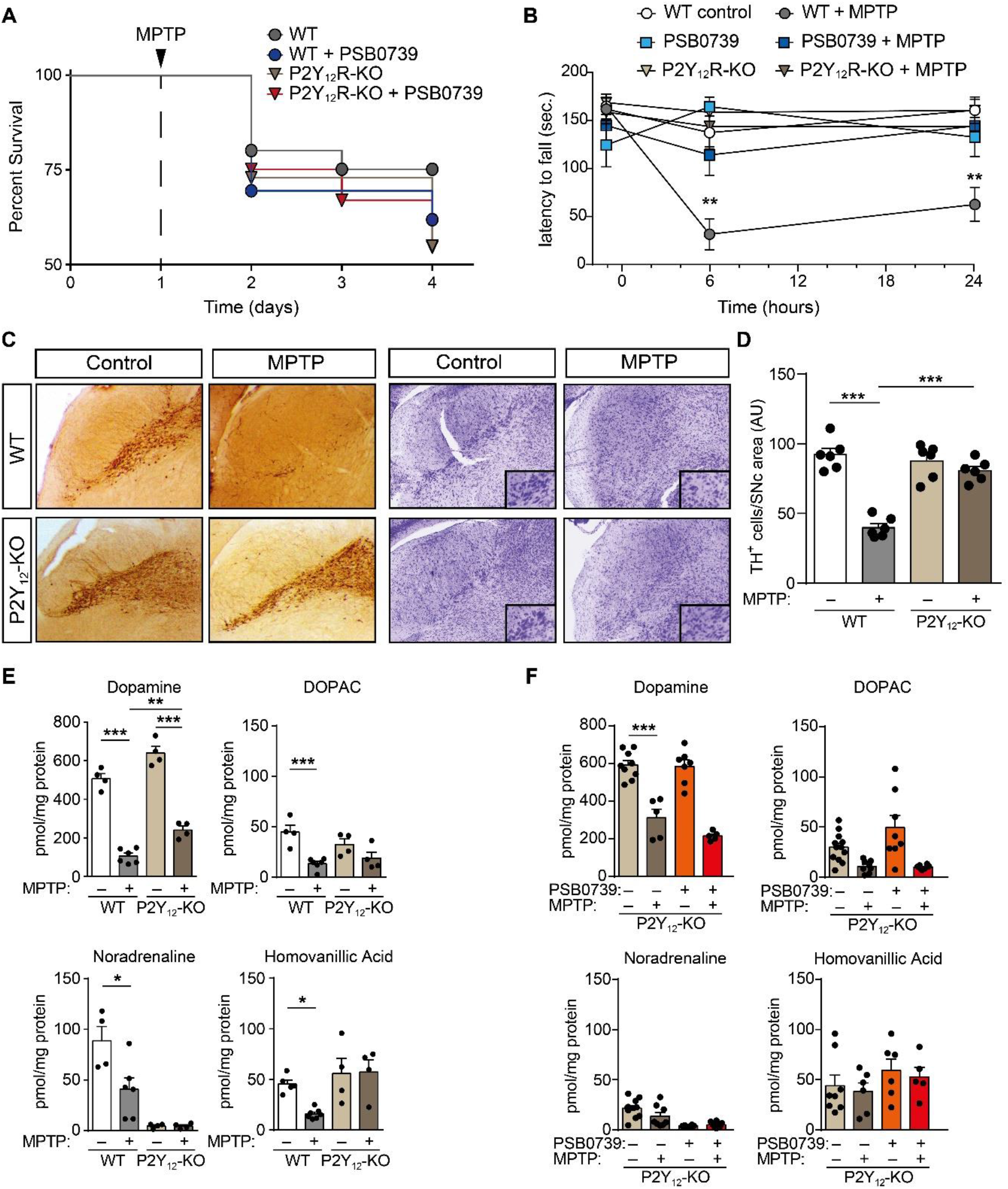
Loss of P2Y_12_R promotes acute mortality but preserves dopaminergic neurons and alleviates PD symptoms after MPTP-treatment. **(A-F)** WT or P2Y_12_-KO mice were pretreated with 0.3 mg / kg PSB 0739 or its vehicle and 4 × 20 mg / kg MPTP or its vehicle as indicated. Kaplan-Meier survival analysis performed on experimental groups receiving MPTP treatment (n=11-24) (A). Effect of P2Y_12_R gene deficiency or 0.3 mg / kg PSB 0739 on the motor performance during MPTP-induced PD measured on the rotarod test (n=5-8) (B). Immuno-DAB staining for TH or cresyl-violet staining on representative tissue sections; immunoreactivity is seen in the cell body and processes of dopaminergic and noradrenergic neurons (C); quantification of TH-positive cells in the substantia nigra pars compacta (n=6) (D). Concentration of dopamine, DOPAC, noradrenaline and homovanillic acid were determined from striatum samples of WT and P2Y_12_-KO mice 72 hours after last MPTP administration (n=4-6) (E). Concentration of dopamine, DOPAC, noradrenaline and homovanillic acid were determined from striatum samples of P2Y_12_-KO mice pretreated with 0.3 mg / kg PSB 0739 or its vehicle, 72 hours after last MPTP administration (n=5-12) (F). Data represent the mean ± SEM; *, *p* ≤ 0.05; **, *p* ≤ 0.01; ***, *p* ≤ 0.001 (two-way ANOVA, with Bonferroni’s *post-hoc* test (B, D, E, F)).

Next, we examined how genetic deletion or pharmacological blockade of P2Y_12_R influences MPTP-induced neurodegeneration and motor function impairment. Administration of MPTP to wild-type mice resulted in severe motor function deterioration, when the latency to fall from rotarod apparatus was measured (Figure 1B). Surprisingly, inhibition of P2Y_12_R, either genetically or with pharmacological blockade ameliorated motor function impairment in MPTP-treated mice (Figure 1B). As expected, we found that administration of MPTP to wild-type mice induced selective degeneration of dopaminergic neurons in the substantia nigra *pars compacta* (Figure 1C and D), demonstrated by reduced tyrosine-hydroxylase immunostaining and the presence of degenerating neurons with condensed nuclei, shrunken cytoplasm and prominent nuclear alterations. MPTP-induced neurodegeneration was associated with reduced levels of monoamines, specifically dopamine, DOPAC, homovanillic acid and noradrenaline (Figure 1E and F, and Supplemental Table 1) and decreased ATP levels in the striatum indicating a state of an energy deficit, as reflected by lower ATP/ADP ratio (Supplemental Figure 1A and B). Genetic deletion of the receptor also protected from dopaminergic neuronal cell death in the substantia nigra *pars compacta* following MPTP-treatment (Figure 1C and D), mitigated the decrease in ATP concentration and alleviated the reduction in MPTP-induced dopamine levels, but had no effect on noradrenaline, DOPAC and homovanillic acid concentration in the striatum (Figure 1E and F, and Supplemental Figure 1A). Additionally, we determined the effect of PSB 0739 on MPTP-induced parkinsonism in P2Y_12_R-KO mice (25). Considering that the inhibitor is unable to cross the blood-brain barrier, we have administered PSB 0739 intrathecally, which ensured that solely centrally expressed receptors are blocked. Pharmacological blockade of P2Y_12_R with PSB 0739 in the receptor knock-out mice had no further effect on monoamine concentration in the striatum following MPTP-treatment, when compared to vehicle treated littermate controls (Figure 1F and Supplemental Figure 1A); indicating that PSB 0739 selectively act on P2Y_12_R, and that the inhibitor requires the presence of functional receptors.

### Pharmacologic inhibition of P2Y_12_-receptor is protective against MPTP-induced neuronal and monoamine loss and reduces neuroinflammation

Following, we investigated the influence of PSB 0739 on MPTP-induced neuroinflammation and microglia activation. MPTP administration promoted the degradation of dopaminergic neurons in the substantia nigra *pars compacta* (Figure 2A and B), and reduced the concentration of monoamines, specifically dopamine, DOPAC, homovanillic acid and noradrenaline (Figure 2C and Supplemental Table 1). Furthermore, treatment with MPTP was also associated with neuroinflammation, characterized by increased levels of TNFα, IL-1β and IL-6 and the anti-inflammatory cytokine IL-10 (Figure 2D and E). Similarly to the constitutive loss of P2Y_12_R, the pharmacological blockade of the receptor protected against MPTP-induced Parkinson’s-like disease; PSB 0739 treatment 18 hours prior to MPTP administration reduced dopaminergic cell loss in the substantia nigra *pars compacta* (Figure 2A and B), attenuated the decrease in dopamine concentration measured in the striatum (Figure 2C), and abolished the increase in cytokine levels in the substantia nigra and the striatum (Figure 2D and E, respectively).

**Figure 2.**
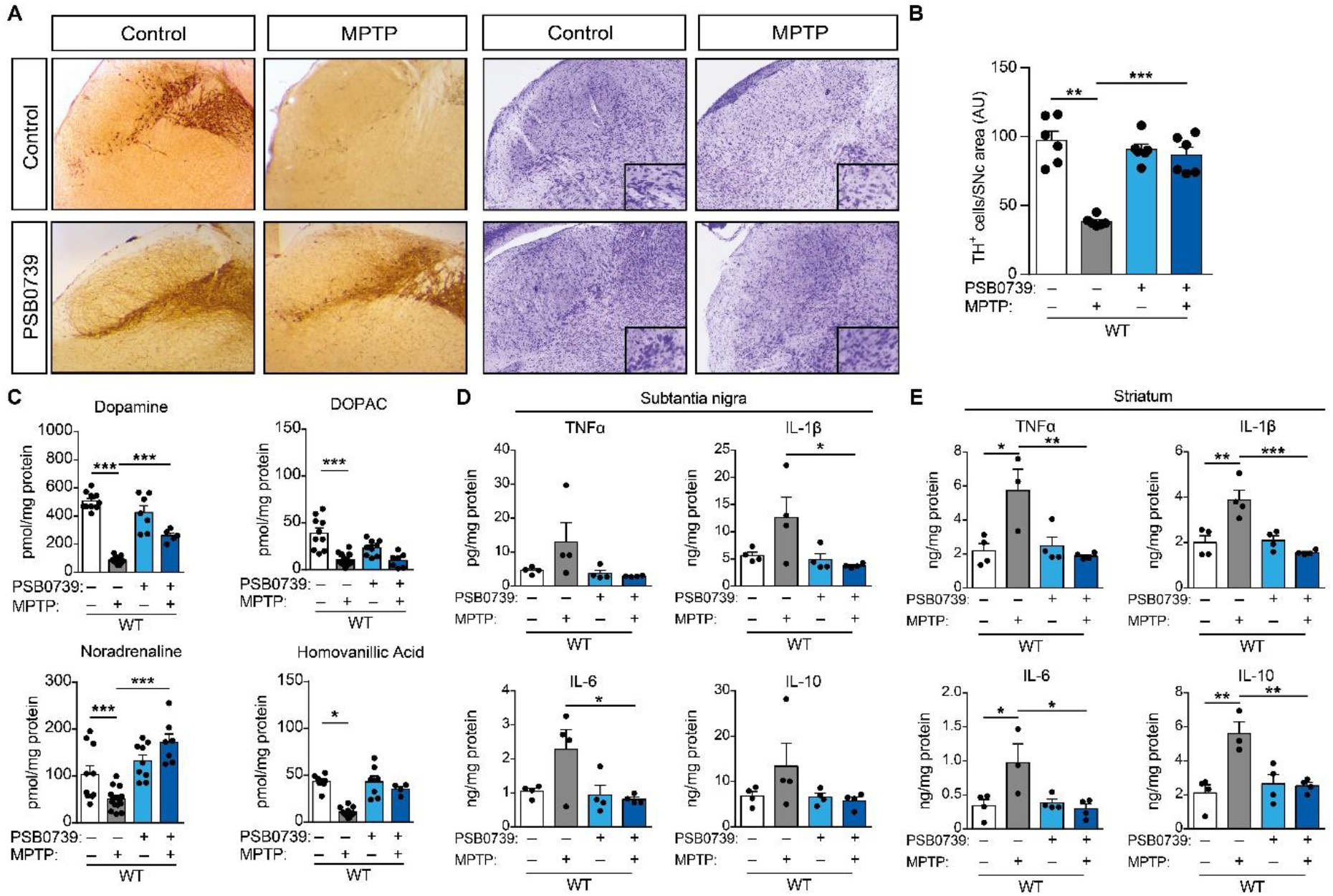
Pharmacological P2Y_12_R inhibition protects against MPTP-induced PD. **(A-E)** WT mice were pretreated with 0.3 mg / kg PSB 0739 or its vehicle and 4 × 20 mg / kg MPTP or its vehicle as indicated. Immuno-DAB staining for TH or cresyl-violet staining on representative tissue sections; immunoreactivity is seen in the cell body and processes of dopaminergic and noradrenergic neurons (A); quantification of TH-positive cells in the substantia nigra pars compacta (n=5-6) (B). Concentration of dopamine, DOPAC, noradrenaline and homovanillic acid were determined from striatum samples 72 hours after last MPTP administration (n=5-12) (C). TNFα, IL-1β, IL-6 and IL-10 concentration measured from substantia nigra (n=4) (D) or striatum samples (n=3-4) (E). Data represent the mean ± SEM; *, *p* ≤ 0.05; **, *p* ≤ 0.01; ***, *p* ≤ 0.001 (two-way ANOVA, with Bonferroni’s *post-hoc* test (B-E)).

These data indicate that P2Y_12_R in the central nervous system has a biphasic effect in the MPTP-induced Parkinson’s disease model. Initially, functional P2Y_12_R are essential to reduce the effects of acute neurotoxicity, where P2Y_12_R–mediated signalling is required to facilitate the recruitment of phagocytic activity of microglia (22, 26); however, during chronic neuronal loss, P2Y_12_R may further contribute to neuroinflammation. Apparently, prolonged activation of P2Y_12_R contributes to maintaining the progression of neurodegeneration, since either pharmacological or genetic inhibition of the receptor improves motor function and protects against dopaminergic cell death in MPTP-induced Parkinson’s disease model; although the marked difference in survival rate may potentially mask the exact function of the receptor in these experimental settings.

To further understand the mechanisms involved, we investigated the involvement of P2Y_12_R on dopamine release from the striatum. After loading the striatal slices with the radioligand, [^3^H]DA, we have found that the uptake of radioactivity (Figure 3A) and the basal dopamine efflux (Figure 3B) was indistinguishable in wild-type control and P2Y_12_R-KO mice. Perfusion with the sodium channel activator, veratridine (5 μM), elicited a sharp and transient elevation in the efflux of [^3^H]DA in wild-type mice, which was reversible upon washout (Figure 3C). The veratridine-evoked [^3^H]DA efflux was higher in the slices of P2Y_12_R-KO mice (Figure 3C and D). A second veratridine challenge elicited a smaller increase under control conditions (Figure 3E), while administration of PSB 0739 15 min before S2 slightly increased the S2/S1 ratio, although this effect did not reach the level of significance (Figure 3E and F). These findings imply that P2Y_12_R gene deficiency modifies dopamine release from nerve terminals; suggesting that in correlation with regulating microglial activity, neuromodulation by P2Y_12_R may also influence extracellular dopamine concentration during Parkinson’s disease.

**Figure 3.**
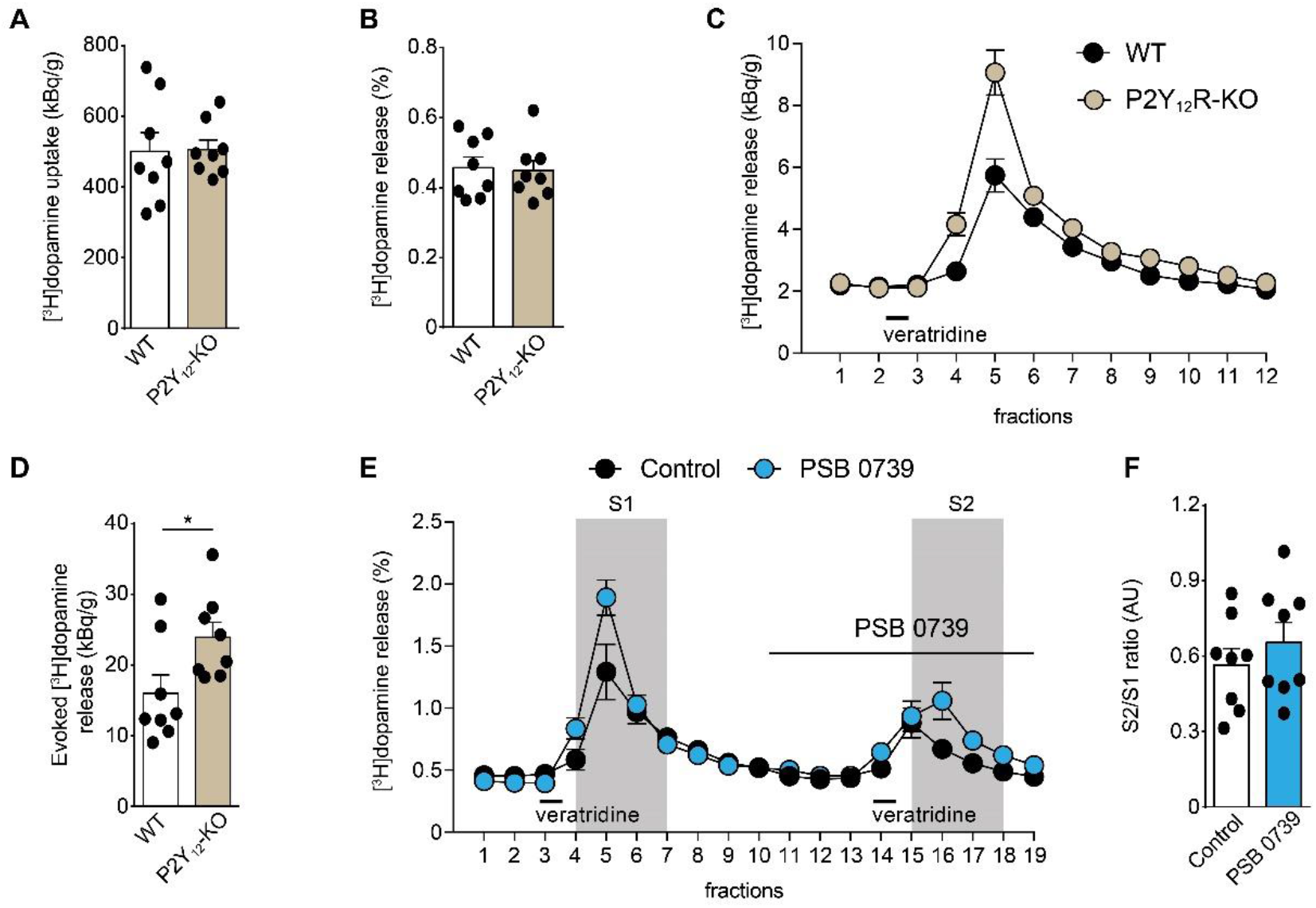
Neuromodulation during P2Y_12_R blockade. **(A-D)** Brain slices were isolated from wild-type and P2Y_12_R-KO mice, and loaded with [^3^H]dopamine. Perfusate samples (fractions) were collected over 3 min periods and assayed for tritium content. Concentration of [^3^H]dopamine uptake were determined from striatum samples (n=8) (A). Resting [^3^H]dopamine efflux from WT and P2Y_12_R-KO mice (n=8) (B). Effect of 5 μM veratridine on [^3^H]dopamine release (n=8) (C); quantification of [^3^H]dopamine release from the striatum (n=8) (D). **(E-F)** Two consecutive veratridine stimuli were applied, and the P2Y_12_R antagonist, PSB 0739 (1 μM, 15 min) was perfused before the second veratridine administration (S2), and [^3^H]dopamine release was measured (n=8) (E); the effect of PSB 0739 on veratridine-evoked [^3^H]dopamine release was expressed as the ratio of S2 over S1, compared to control S2/S1 ratio (n=8) (F). Data represent the mean ± SEM; *, *p* ≤ 0.05 (unpaired two-tailed Student’s *t* test (D)).

### P2Y_12_-receptor blockade halts disease progression after subchronic MPTP administration

Since our data implicate a pivotal role for P2Y_12_R in neuroinflammation, we further investigated the contribution of P2Y_12_R during subchronic neurodegeneration. To account for the markedly different survival rate of wild-type and P2Y_12_R deficient animals during the acute experimental Parkinson’s disease model, an alternative MPTP treatment protocol was used to explore P2Y_12_R actions during disease progression; rather than administering the P2Y_12_R-inhibitor prior to MPTP, a subchronic Parkinson’s disease model was generated by a single daily administration of MPTP over the course of five days, followed by receptor blockade for four consecutive days. During the early phase of the MPTP treatment, on day five to six, limited effect could be observed on dopaminergic neuron loss in wild-type and P2Y_12_R-KO mice (Figure 4A and B). However, twenty-one days following MPTP administration, we detected significant loss of dopaminergic neurons in the substantia nigra *pars compacta* (Figure 4C and D), reduction in dopamine, DOPAC, noradrenaline and homovanillic acid levels (Figure 4E), and markedly decreased motor functions (Figure 4F). P2Y_12_R inhibition increased dopaminergic cell survival (Figure 4C and D), reduced the decrease in dopamine levels (Figure 4E), and improved motor function (Figure 4F) assessed three weeks after treatment. Furthermore, using this model, there was no difference in the survival rate between the experimental groups (Figure 4G).

**Figure 4.**
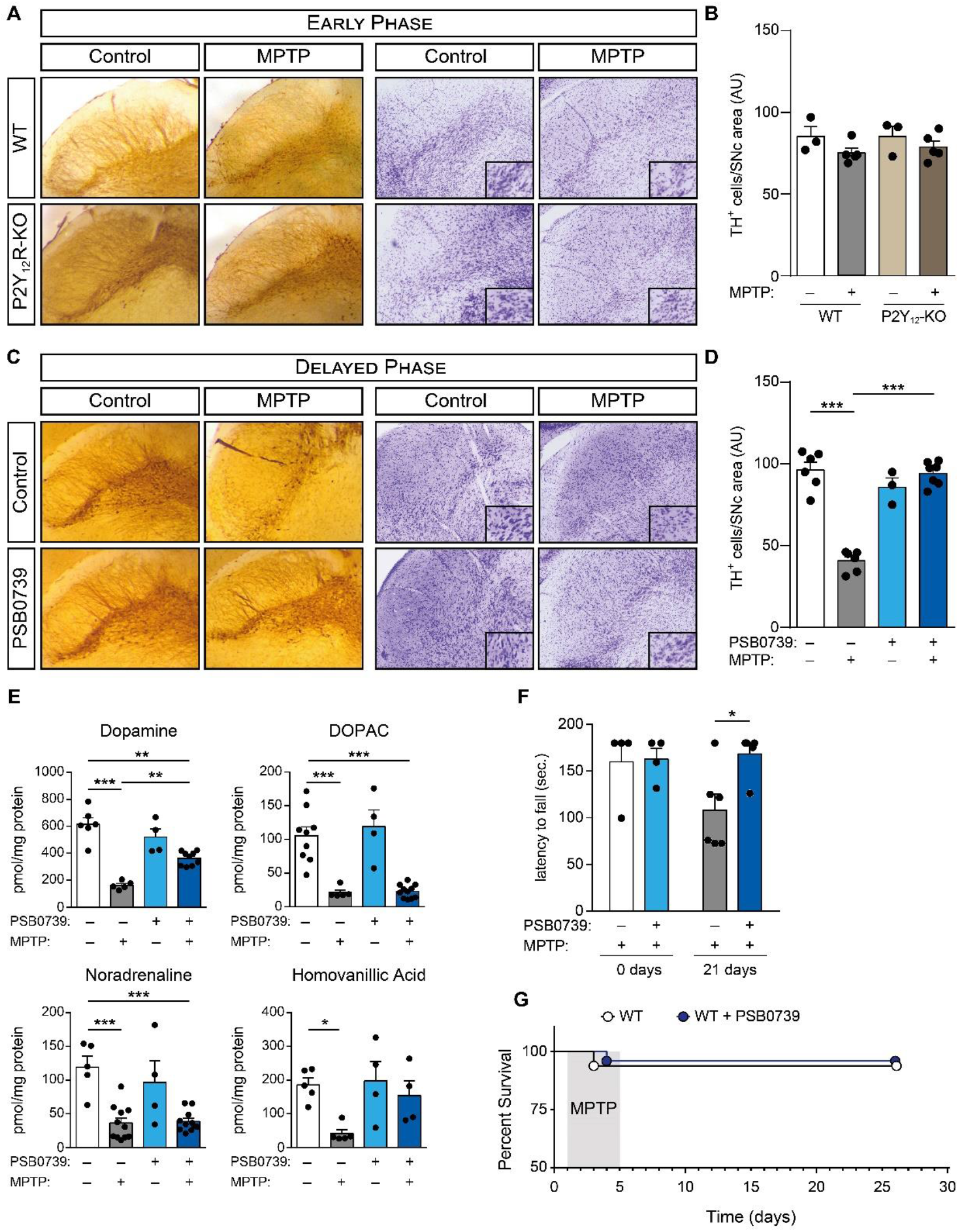
P2Y_12_R blockade halts disease progression. **(A-G)** WT or P2Y_12_-KO mice were treated with 20 mg / kg MPTP daily for five consecutive days, followed by treatment with 0.3 mg / kg PSB 0739 or its vehicle. Immuno-DAB staining for TH and cresyl-violet staining on representative tissue sections five days after the initial MPTP administration; immunoreactivity is seen in the cell body and processes of dopaminergic and noradrenergic neurons (A); quantification of TH-positive cells in the substantia nigra pars compacta (n=3-5) (B). Immuno-DAB staining for TH and cresyl-violet staining on representative tissue sections twenty-one days after the last MPTP administration (C); quantification of TH-positive cells in the substantia nigra pars compacta (n=3-7) (D). Concentration of dopamine, DOPAC, noradrenaline and homovanillic acid were determined from striatum samples 21 days after last MPTP administration (n=4-12) (E). Effect of PSB 0739 or its vehicle on the motor performance during MPTP-induced PD measured on the rotarod test before and 21 days after treatment (n=4-6) (F). Kaplan-Meier survival analysis performed on experimental groups receiving MPTP treatment (n=8-16) (G). Data represent the mean ± SEM; *, *p* ≤ 0.05; **, *p* ≤ 0.01; ***, *p* ≤ 0.001 (two-way ANOVA, with Bonferroni’s *post-hoc* test (B-F)).

### P2Y_12_-receptor mediates microglia activation via Rho-kinase and p38 MAPK phosphorylation

To characterize the intracellular mechanism underlying the protective effect of P2Y_12_R blockade, the relationship between P2Y_12_R stimulation and microglia activation was investigated. Initially, since the activation of p38 mitogen-activated protein kinase (p38 MAPK) can directly promote or indirectly stimulate cytokine production via MK2 or MSK1/2 in microglia (27–30), changes in the p38 MAPK activity was measured. Phosphorylation of p38 MAPK at Threonine 180 / Tyrosine 182 increases the kinase activity, thus promoting phosphorylation of several transcriptional factors and further bolster inflammatory cytokine biosynthesis (31). In initial experiments we confirmed that treatment with LPS (1 μg / mL, 60 min) or the P2Y_12_R agonist ADP (10 μM, 60 min) promotes p38 MAPK phosphorylation at Threonine 180 / Tyrosine 182 *in vitro* (Figure 5A and B) using murine microglial cell line BV-2. The small GTPase, Ras homolog family member A (RhoA) and the downstream effector, Rho-associated, coiled-coil containing protein kinase 1 (ROCK1) has been previously shown to mediate p38 MAPK phosphorylation and activation (32); we have tested the effect of the ROCK1 inhibitor, Y-27632, on LPS or ADP-induced p38 MAPK activation. Pretreatment with Y-27632 (10 μM, 30 min) markedly decreased LPS or ADP-induced p38 MAPK phosphorylation (Figure 5B); additionally, pretreatment with the selective P2Y_12_R inhibitor, PSB 0739 (500 nM, 30 min) blocked ADP-induced, but not LPS-induced p38 MAPK activation (Figure 5A). These results would indicate that under *in vitro* conditions, both P2Y_12_R and ROCK1 are necessary for ADP-mediated p38 MAPK phosphorylation.

**Figure 5.**
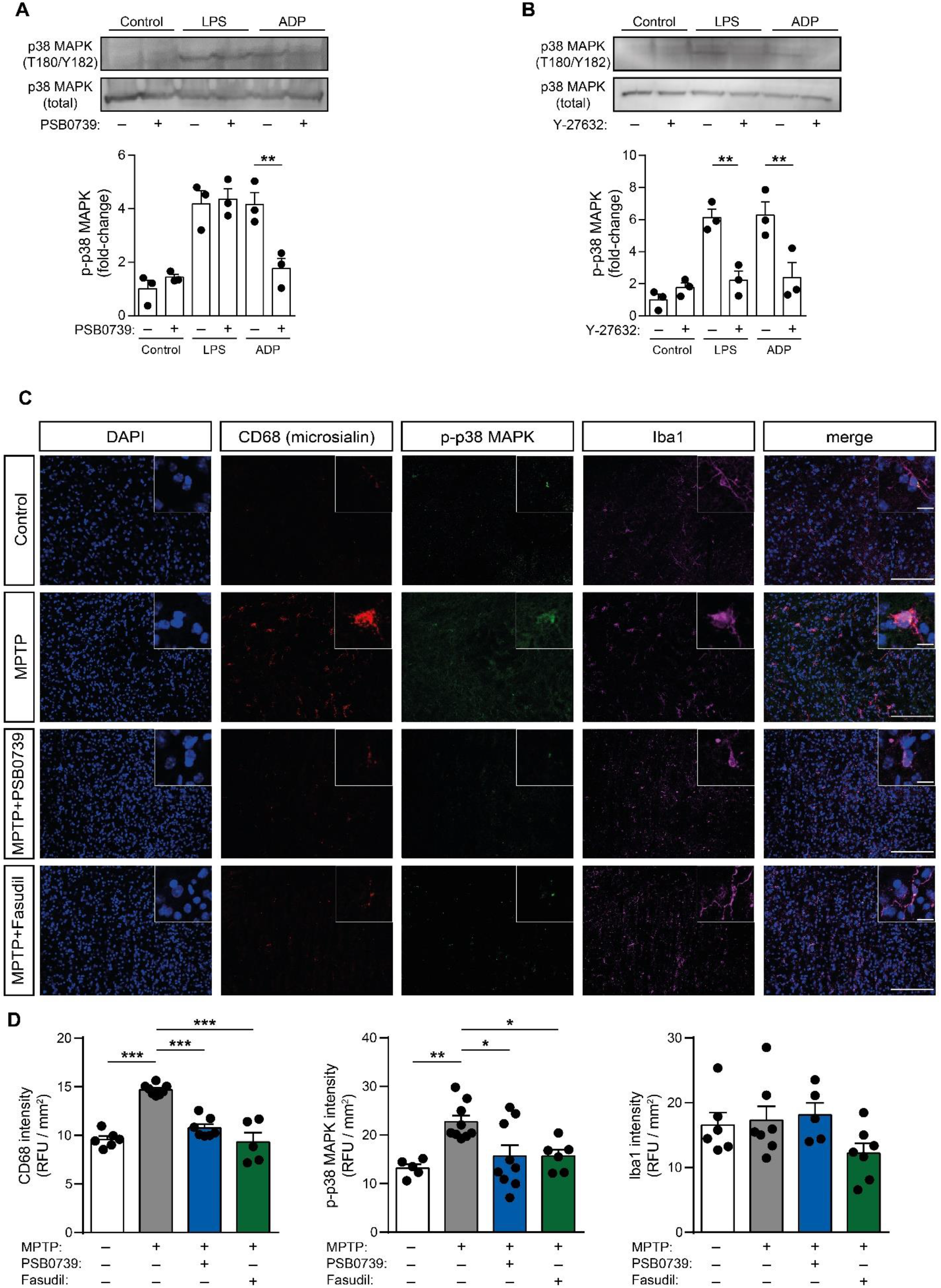
P2Y_12_R mediates MPTP-induced microglia activation and controls Rho-kinase-dependent p38 MAPK phosphorylation. **(A-B)** Murine BV2 microglia cells were pretreated with PSB 0739 (500 nM, 30 min) (A) or with Y-27632 (10 μM, 30 min) (B) and were incubated with LPS (100 ng / mL, 60 min), ADP (10 μM, 5 min) or solvent (control) and phosphorylation of p38 MAPK at Threonine 180 / Tyrosine 182 were determined by immunoblotting. Graphs show the densitometric evaluation (n=3). **(C)** WT mice were treated with 20 mg / kg MPTP daily for five consecutive days, followed by treatment with 0.3 mg / kg PSB 0739, 50 mg / kg body weight per day fasudil or its vehicle. Shown are representative immuno-confocal microscopy images of brain slices isolated from WT mice stained with antibodies directed against CD68 (microsialin, red), phosphorylated-p38 MAPK (green), Iba1 (purple), DAPI (blue) and overlay image (merge). Scale bar: 100 μm, corresponds to 20 μm inset **(D)** Quantification of CD68 fluorescence intensity (left panel), phosphorylated-p38 MAPK fluorescence intensity (middle panel) and Iba1 fluorescence intensity (right panel) (n=5-7, left panel; n= 5-9, middle panel; n= 5-7, right panel). Data represent the mean ± SEM; *, *p* ≤ 0.05; **, *p* ≤ 0.01; ***, *p* ≤ 0.001 (two-way ANOVA with Bonferroni’s *post-hoc* test (A, B, D)).

Next, the involvement of P2Y_12_R and ROCK1 blockade on p38 MAPK phosphorylation and microglia activation under *in vivo* conditions was tested. The myeloid specific lysosomal-associated membrane protein, CD68, was characterized as a microglial activation marker previously (33, 34); while the ionized calcium-binding adapter molecule 1 (Iba1) is known to be strongly and specifically expressed in both senescent and activated microglia (35). In our experiments, MPTP administration markedly increased phospho-p38 MAPK T180/Y182 and CD68 fluorescence intensity in brain slices in the substantia nigra area compared to saline treated control animals, whereas Iba1 intensity was unchanged (Figure 5C and D). Intrathecal administration of 0.3 mg / kg PSB 0739 alone was without effect (data not shown). However, P2Y_12_-receptor blockade significantly reduced MPTP-induced p38 MAPK activity and microglia activation, measured by phospho-p38 MAPK T180/Y182 and CD68 positivity (Figure 5C and D). To assess ROCK1 involvement, the specific ROCK1 inhibitor fasudil, was orally administered in the drinking water (50 mg / kg body weight per day) (36). Fasudil-treatment alone had no effect on phospho-p38 MAPK T180/Y182 or CD68 fluorescence intensity (data not shown). When fasudil was administered during MPTP-treatment, phospho-p38 MAPK T180/Y182 and CD68 fluorescence intensity was significantly reduced in comparison to MPTP-treatment alone (Figure 5C and D). No changes could be observed in both p38 MAPK and Iba1 fluorescence intensity between control, MPTP treated or MPTP combined with PSB 0739 or fasudil-treated groups (Figure 5C and D; and Supplemental Figure 2A and B).

Since our data strongly suggest ROCK1 to be also involved in MPTP-induced neuroinflammation, we further investigated the involvement of ROCK1 during neurodegeneration. Similar to our previously shown results utilizing the subchronic Parkinson’s disease model, MPTP administration induced significant loss of dopaminergic neurons in the substantia nigra *pars compacta* (Figure 6A and B), reduction in dopamine, DOPAC, noradrenaline and homovanillic acid levels (Figure 6C), and markedly decreased motor functions (Figure 6D). Fasudil-treatment increased dopaminergic cell survival (Figure 6A and B) and improved motor function (Figure 6D). Interestingly, treatment with fasudil only (without MPTP administration) substantially reduced monoamine levels in the striatum (Figure 6C); however, the decrease in dopamine concentration compared to the control conditions did not reach statistical significance. When fasudil-treatment was combined with subchronic MPTP administration, the decrease in dopamine concentration in the striatum was alleviated compared to the MPTP-induced decrease (Figure 6C). Apparently, the Rho-kinase inhibitor fasudil, while evidently have a protective effect considering dopaminergic neuron loss in addition to preserving motor function during experimental Parkinsonism; prolonged administration of the drug leads to a marked decrease in striatal monoamine levels. Furthermore, we could not find difference in the survival rate in response to fasudil administration in MPTP treated mice (Figure 6E).

**Figure 6.**
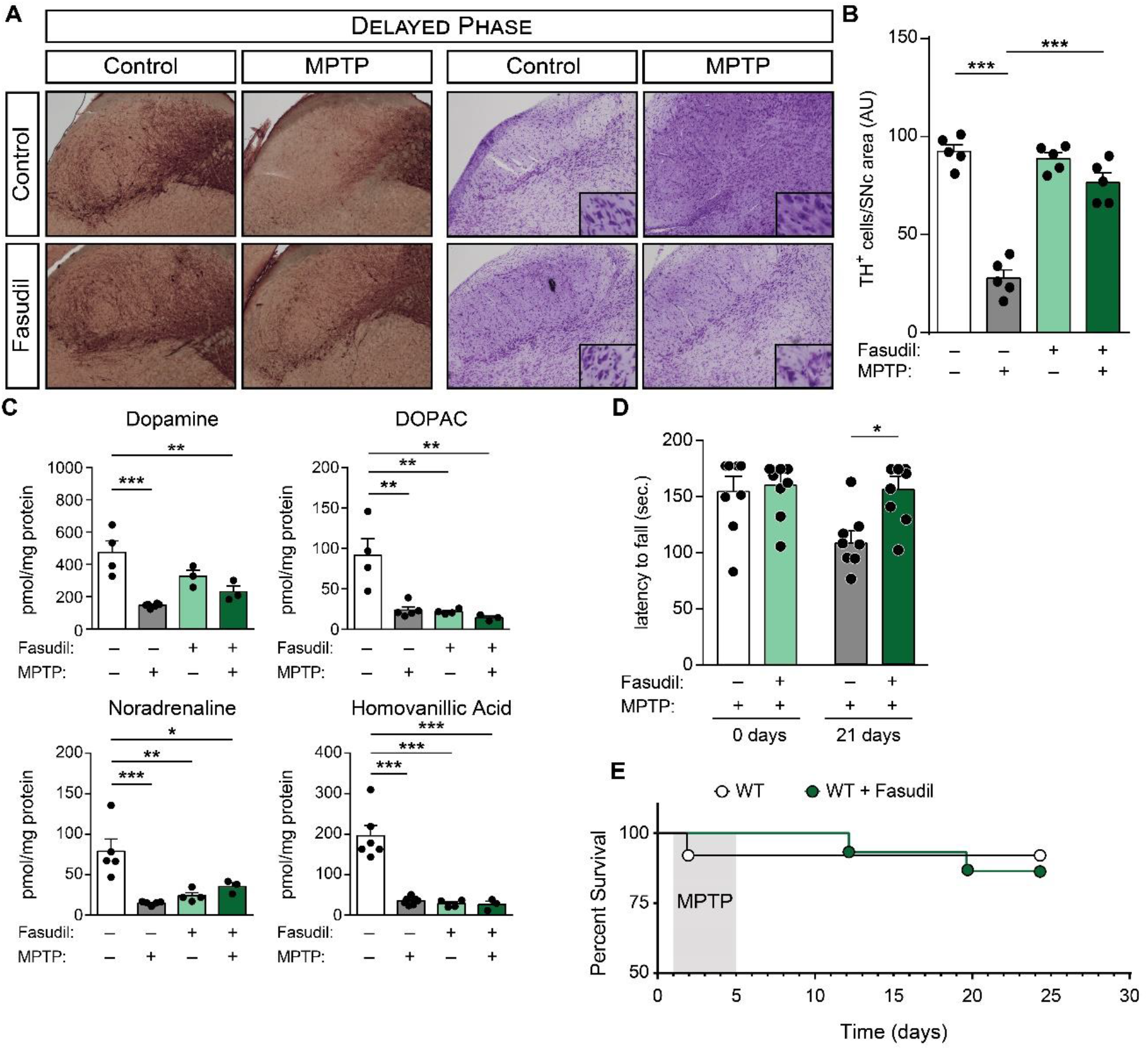
Pharmacological Rho-kinase blockade interrupts disease progression and alleviates MPTP-induced parkinsonism. **(A-E)** WT mice were treated with 20 mg / kg MPTP daily for five consecutive days, followed by treatment with 50 mg / kg body weight per day fasudil or its vehicle. Shown are representative images of immuno-DAB staining for TH and cresyl-violet staining on tissue sections twenty-one days after the initial MPTP administration; immunoreactivity is seen in the cell body and processes of dopaminergic and noradrenergic neurons (A); quantification of TH-positive cells in the substantia nigra pars compacta (n=5) (B). Concentration of dopamine, DOPAC, noradrenaline and homovanillic acid were determined from striatum samples 21 days after last MPTP administration (n=3-8) (C). Effect of fasudil or its vehicle on the motor performance during MPTP-induced PD measured on the rotarod test before and 21 days after treatment (n=8-9) (D). Kaplan-Meier survival analysis performed on experimental groups receiving MPTP treatment (n=8-10) (E). Data represent the mean ± SEM; *, *p* ≤ 0.05; **, *p* ≤ 0.01; ***, *p* ≤ 0.001 (two-way ANOVA, with Bonferroni’s *post-hoc* test (B-D)).

Lastly, no changes could be observed in P2Y_12_R fluorescence intensity between control, MPTP treated or MPTP combined with PSB 0739 or Fasudil-treated groups (Supplemental Figure 3A and B). The specificity of the P2Y_12_R antibody has been validated using wild-type control and P2Y_12_R-KO mice (Supplemental Figure 4A and B).

### P2Y_12_-receptor is expressed on activated human microglia in Parkinson’s disease associated dementia

The relevance of our findings is supported by marked expression of P2Y_12_R on cells with typical morphology of activated microglial cells in the neocortex (Figure 7A and B) and striatum (Figure 7C) of patients with Parkinson’s disease dementia (i.e. Parkinson’s disease patients who developed dementia at the late stage of the disease); although to lesser extent than in Alzheimer’s disease (Figure 7D, E and F), which is usually associated with more severe neuronal loss and abnormal protein burden.

**Figure 7.**
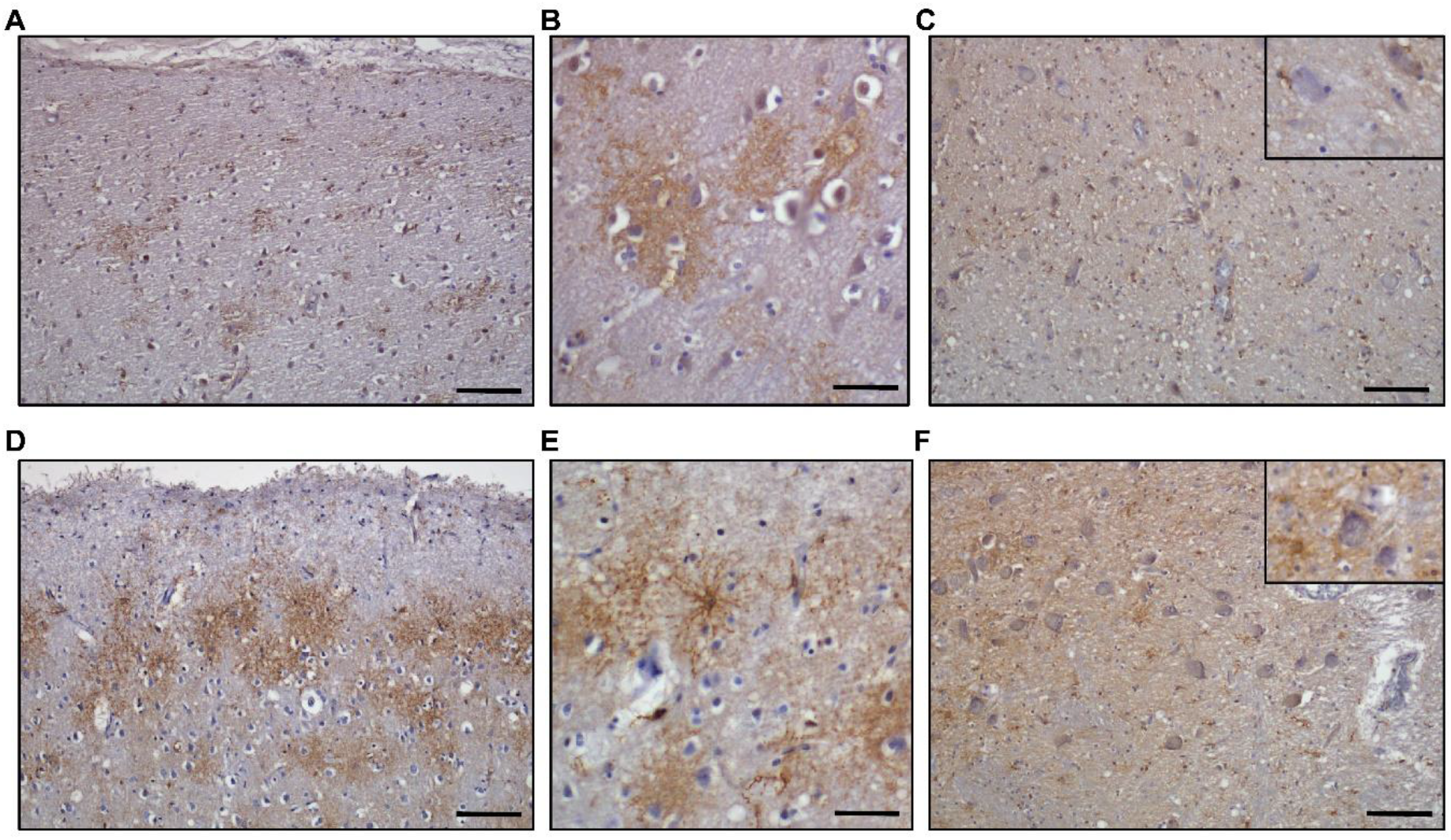
Human P2Y_12_R expression in neurodegenerative diseases. **(A-C)** Representative images from human frontal lobe neocortex samples (A,B) or human striatum sample from patients with Parkinson’s Disease Dementia (C). Immunohistochemistry was performed against human P2Y_12_R and visualized with DAB chromogen and haematoxylin counterstain. (Scale bar: 100 μm (A,C), corresponds to 50 μm in B and inset (C)) **(D-F)** Representative images from human frontal lobe neocortex samples (D,E) or human striatum sample from patients with Alzheimer’s Disease (F). Immunohistochemistry was performed against human P2Y_12_R and visualized with DAB chromogen and haematoxylin counterstain. (Scale bar: 100 μm (D,F), corresponds to 50 μm in E and inset (F)).

## Discussion

Neuroinflammation has a central role in Parkinson’s disease pathogenesis (8, 10, 37); characterized by microglial proliferation, activation and production of inflammatory mediators (37, 38). Various studies reported on compounds affecting neuroinflammation and ameliorating PD symptoms; most recently, a novel Krf2 activator, KKPA4026, has been shown to attenuate motor deficit associated with PD and exhibit protective effect on dopaminergic neurons (39). Furthermore, inhibition of NLRP3 inflammasome by dimethylaminomylide alleviates neuroinflammation in MPTP-induced PD model (40). Administration of vitamin D for 10 consecutive days proved beneficial in PD animal model (41); whereas modulating microglia proliferation via inhibition of CSF1R by GSW2580 markedly reduced mRNA levels of pro-inflammatory mediators and mitigated the loss of dopaminergic neurons and motor deficits during PD (42). While microglia seemingly maintain neuroinflammation and exacerbate Parkinson’s disease symptoms, the presence of functional microglia also appears to be essential to reduce neurotoxic effects, which issue has been highly controversial in PD and in other forms of neurodegeneration. For example, it has been shown that microglia depletion via the CSF1R inhibitor PLX3397, aggravates the impairment in locomotor activity, worsens the loss of dopaminergic neurons and augments the infiltration of leukocytes into the brain after MPTP treatment (24).

Here we provide novel insights into the mechanisms of experimental PD to resolve some of the controversies presented above. In addition, we report for the first time the involvement of P2Y_12_ receptors in experimental PD. P2Y_12_-receptors are preferentially expressed on microglia in the CNS (43), which makes it likely that the primary target for intrathecally administered PSB 0739 are microglia. Since P2Y_12_-KO and PSB 0739-treated mice displayed similar effects on disease progression our results suggest that while the presence of P2Y_12_R is essential to prevent acute neurotoxicity and associated mortality, prolonged activation of P2Y_12_R may alter pro- and anti-inflammatory cytokine production and maintain neuroinflammation during experimental Parkinson’s disease. Chronic inhibition of P2Y_12_-receptors during disease progression mitigates inflammation, protects against dopaminergic neuronal cell loss and alleviates motor impairments. Our results indicate that the presence of P2Y_12_-receptors in the CNS is essential during the initiating phase of the experimentally induced Parkinson’s disease model. This is likely to be mediated by the interactions between P2Y_12_R-positive microglia and neurons. However, P2Y_12_R also contributes to phagocytic activity of microglia (20, 22, 26) and shapes cytokine production (33, 44) presumably by the reduction of p38 MAPK phosphorylation via RhoA-ROCK1 dependent mechanism. Besides regulating microglial activity, neuromodulation by P2Y_12_R might also affect extracellular dopamine concentration during Parkinson’s disease, since our [^3^H]dopamine release experiments indicate that P2Y_12_R gene deficiency modulate transmitter release from nerve terminals; however, pharmacological blockade of the receptor had limited influence on dopamine release. While the involvement of neuromodulation during MPTP-induced experimental parkinsonism via P2Y_12_R may suggest an important mechanism to adjust dopamine concentration in the striatum, to unmistakably address the role of neuromodulation during PD requires further experiments.

Recently, the potentially advantageous effect of Rho-kinase inhibition in several neurodegenerative disorders has been demonstrated (26, 45). Fasudil-administration proved to inhibit axonal degeneration and neuronal death in an experimental Parkinson’s disease model, when administered prior to disease onset (46, 47).

Furthermore, ROCK inhibitors were postulated to promote the activity of Parkin-mediated mitophagy pathway during Parkinson’s disease, by recruiting HK2, a positive regulator of Parkin, to mitochondria. The increased Parkin recruitment to damaged mitochondria leads to increased removal of mitochondria from injured cells (45). However, a side effect of Rho-kinase inhibition, namely the stimulation of monoamine oxidase activity, was previously demonstrated (48). Since increased monoamine oxidase activity leads to rapid metabolism of monoamines, this crucial observation may also explain our finding that prolonged fasudil-treatment lead to the reduction in central monoamine levels, and proposes a potential limitation on the application of fasudil in the treatment of Parkinson’s disease. Our finding that P2Y_12_R stimulation regulates ROCK1 activity and consequently p38 MAPK-dependent cytokine production furthers our understanding of the intracellular mechanistic pathway during Parkinson’s disease; while the inhibition of Rho-kinase activity determine the clearance of defective mitochondria and govern cytokine signalling to influence their immediate surroundings, it also affects monoamine oxidase activity and reduces central monoamine concentration. In comparison, modulating P2Y_12_R activity has a highly specific protective influence in MPTP-induced parkinsonism, without the observed adverse effects of fasudil; consequently P2Y_12_R-inhibition might be more advantageous in the treatment of Parkinson’s disease.

Recent work performed by Andersen et al. (49), using summary statistics from recent meta-analysis genome-wide association studies in Parkinson’s disease patients, identified significant enrichment of Parkinson’s disease risk heritability in microglia and monocytes. Furthermore, *P2RY12* was indicated as the strongest candidate gene driving a significant Parkinson’s disease association signal. Their finding, while lacking data, whether functionally relevant polymorphisms are identified in the proposed *P2RY12* candidate gene region; nevertheless corroborates our finding that neuroimmune mechanisms during PD pathogenesis as well as the apparent microglia dysregulation are strongly influenced by P2Y_12_R function in human patients.

P2 purinergic receptors are nucleotide sensitive receptors, which are involved in both the maintenance of neuroinflammation and facilitation of tissue repair (50, 51). Microglia have been shown to express ionotropic (P2X4 and P2X7) and metabotropic (P2Y1, P2Y2, P2Y6 and P2Y_12_) purinergic receptors (20, 52, 53), which are activated by the released nucleotides during inflammation. Previous work has already shown P2Y_12_ and P2X_4_ receptors to regulate microglial activation, migration and polarization via phosphoinositde-3-kinase (PI3K) activation and Akt phosphorylation (20, 54). Recently, P2Y6-receptor expression has been shown to increase in patients with Parkinson’s disease and was proposed as a potential biomarker in PD (53). In the current study, we also demonstrated that P2Y_12_R are expressed on activated microglia in human patients suffering from neurodegenerative diseases. P2Y_12_R expression was prominent in patients with Alzheimer’s disease, which is associated with severe neuronal loss and abnormal protein burden, and to a lesser degree, in patients with Parkinson’s disease dementia. While P2Y_12_-receptor has been established as an important regulator for microglial activation, its contribution to neurodegeneration in PD has not been explored until now.

P2Y_12_-receptors exhibit unique expression pattern; they are highly expressed on platelets and microglia (55). Pharmacological inhibitors are widely used in clinical practice to target P2Y_12_R in order to inhibit platelet aggregation and thereby prevent stroke and myocardial infarction (56). Furthermore, we previously showed that P2Y_12_R regulate hyperalgesia and local inflammatory processes, and also contribute to the development of chronic inflammation (18); while P2Y_12_R located on microglia affect neuropathic pain via GTP-RhoA/ROCK pathway and p38 MAPK activation after spinal cord injury (57). Recently, we revealed the pivotal role for P2Y_12_R during microglia-neuron communication through somatic purinergic junctions (19); where the interaction sites between the neuronal cell bodies and microglial processes create a highly specialized nanoarchitecture through which microglia monitor neuronal activity and mediate neuroprotection via P2Y_12_R upon acute brain injury. These interactions may explain the early protective actions by microglial P2Y_12_R to prevent mortality after MPTP administration, while these sites may also function as initiators for microglial phagocytosis around terminally injured neurons (19).

Microglia, the tissue-specific macrophages of the CNS perform constant surveillance of the brain parenchyma under physiological circumstances. They exhibit ramified morphology, where long cytoplasmic processes survey their surrounding environment (58). Exogenous stimuli, such as bacterial endotoxin lipopolysaccharide (LPS), IFNγ, trauma, reperfusion-injury or chemical toxins induce microglial activation (59) to stimulate the production of iNOS, modulate the expression of various cell surface markers, like CD68, CD11b and MHC-II, and to increase pro-inflammatory cytokine (IL-1β, IL-6 and TNFα) and chemokine (CCL2, CXCL10) production (60–62). On the other hand, microglia is indispensable for the resolution of inflammation, where microglia produces anti-inflammatory mediators, such as IL-4, IL-10, extracellular matrix proteins or glucocorticoids (38).

We can only speculate about the molecular mechanisms that link P2Y_12_ signalling with cytokine production in the present experimental model. We hypothesise that damage-associated molecular patterns (DAMPs) bind to P2Y_12_-receptors, which via the Gα_i_ subunit reduces the intracellular cAMP concentration. The consequential decrease in protein kinase A activity results in increased RhoA activity, presumably due to the decreased phosphorylation state of the inhibitory Serine 188 site on RhoA, leading to increased ROCK activity (32). Enhanced ROCK activity further promotes Threonine 180 / Tyrosine 182 phosphorylation on p38 MAPK, whereas the simultaneous double phosphorylation on p38 MAPK stimulates the kinase activity inducing the transcription and production of cytokines (63) (Figure 8). Interestingly, aside from ROCK-mediated activation of p38 MAPK, the MAPK pathway can also be activated by cytokines, namely IL-1β and TNFα (64, 65); providing an enhancing feed-back loop to inflammation. This can lead to a circumstance, where a specific stimulus activating p38 MAPK promotes inflammation and cytokine production, and enters into a vicious cycle, where inflammation maintains inflammation. However, if this cycle could be selectively disrupted, disease resolution would be an advantageous outcome.

**Figure 8.**
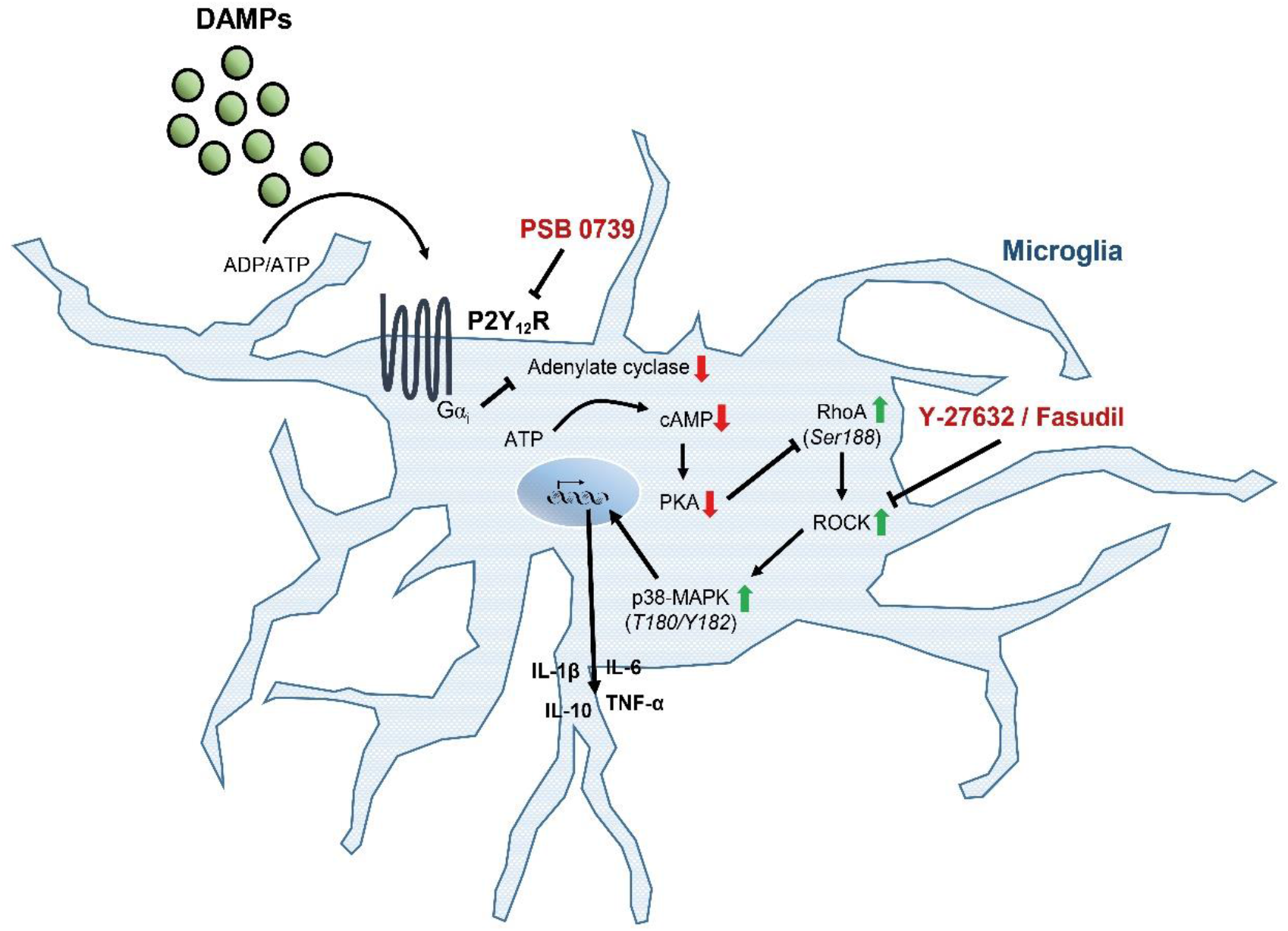
Putative model of microglial P2Y_12_-receptor and its downstream signaling mechanism during disease progression in Parkinson’s disease. Neurodegeneration during Parkinson’s disease causes the release of damage-associated molecular pattern molecules (DAMPs), such as ADP and ATP. The released nucleotides active P2Y_12_-receptor (P2Y_12_R) expressed on microglia, and subsequently through activation of the heterotrimeric G protein Gi results in the inhibitory regulation of adenylate cyclase. Decreased levels of cAMP leads to reduced protein kinase A (PKA) activity, which in turn results in reduced phosphorylation of RhoA at serine 188. Phosphorylation of RhoA at serine 188 negatively regulate RhoA activity; therefore, the decreased phosphorylation state results in increased RhoA and in turn, enhanced ROCK and p38 MAPK activity, which then leads to bolstered pro- and anti-inflammatory cytokine biosynthesis. Arrows indicate stimulation, blunt lines represent inhibition. Red arrows indicate decrease in concentration or activity, green arrows indicate increased enzymatic activity. PSB 0739 is a specific inhibitor of P2Y_12_R, Y-27632 is a specific inhibitor for ROCK.

A fundamental function of microglia is their ability to respond to harmful stimuli and trigger inflammatory response, while also to initiate disease resolution and neuroprotection. We have identified the Gi-coupled P2Y_12_-receptor to sense the increased levels of nucleotides released during cellular damage, and to initiate cytokine production by regulating p38 MAPK activity via ROCK1. Blockade of P2Y_12_R is harmful in the acute phase of MPTP-induced parkinsonism, but prevents MPTP-induced dopaminergic neuron loss and the development of Parkinson’s disease at later time points. Furthermore, inhibition of the receptor abrogates disease progression, reduces motor function impairment and mitigates neuronal cell death. Modulating P2Y_12_R activity appears to be highly selective and specific compared to Rho-kinase inhibition, which has a wide-range of unfavourable side effects; therefore, regulating P2Y_12_R function might be a more favourable target in the treatment of Parkinson’s disease. Thus, understanding the complex mechanisms through which P2Y_12_R signalling in the CNS contributes to injury or repair may allow targeted modulation of microglial responses and neuroinflammation to shape disease progression during Parkinson’s disease.

## Methods

### Reagents

1-methyl-4-phenyl-1,2,3,6-tetrahydropyridine (MPTP, Cat. No. M0896), adenosine 5’-diphosphate (Cat. No. A2754), bovine serum albumin (Cat. No. A2153), lipopolysaccharide (Cat. No. L2880) and veratridine (Cat. No. V5754) were purchased from Sigma-Aldrich. Y-27632 were obtained from Cayman Chemicals (Cat. No. 129830-38-2). PSB 0739 were from Tocris Bio (Cat. No. 3983) and Fasudil-HCl were from LC Laboratories (Cat. No. F-4660). Antibodies directed against phosphorylated p38 MAPK (Cat. No. 9211) and p38 MAPK (Cat. No. 9212) were purchased from Cell Signaling Technology, against tyrosine hydroxylase was from Sigma-Aldrich (Cat. No. AB152), against P2Y_12_-receptor was from AnaSpec (Cat. No. AS-55043A), against Iba1 was from Synaptic System (Cat. No. 234004), against CD68 was purchased from Bio-Rad Laboratories (Cat. No. MCA1957).

### Primary Cells

Murine immortalized microglia cell line (BV2) were cultured in Dulbecco’s modified Eagle’s medium supplemented with heat inactivated FBS (10 %), insulin, non essential amino acid, penicillin-streptomycin and gentamycin at pH 7.2-7.4 (ThermoFisher Scientific, Cat No. 11966025).

### Western blotting

Cells were lysed in radioimmunoprecipitation assay (RIPA) buffer containing 150 mM NaCl, 50 mM Tris-HCl (pH 7.4), 5 mM EDTA, 0.1 % (w/v) SDS, 0.5 % sodium deoxycholate and 1 % Triton X-100 as well as protease inhibitors (10 mg / ml leupeptin, pepstatin A, 4-(2-aminoethyl) benzensulfonyl-fluorid and aprotinin) and phosphatase inhibitors (PhosSTOP−, Roche). Total cell lysates were separated by sodium dodecyl sulfate-polyacrylamide gel electrophoresis. Protein was then transferred onto nitrocellulose membranes, followed by overnight incubation with primary antibodies. Membranes were incubated with horseradish peroxidase-conjugated secondary antibodies (Cell Signaling Technology) for 1 hour room temperature and were developed using the ECL detection system (Thermo Scientific Pierce, Life Technologies). In cases where protein phosphorylation was analyzed using phosphosite-specific antibodies, membranes were first developed with anti-phosphosite-specific antibodies. After evaluation, antibody dissociation from the membrane was induced using Restore™PLUS Western Blot Stripping Buffer (ThermoFisher Scientific) according to the manufacturer’s instructions, and membranes were then reprobed with antibodies recognizing the corresponding protein. Protein band intensities were analyzed by ImageJ software (NIH). Intensity values of bands representing phosphorylated sites of proteins were normalized to the intensity of the band representing total protein.

### Cresyl-violet (Nissl) histochemistry

Mice were sacrificed by CO_2_ inhalation and the chest cavity was opened for perfusion with 4 °C phosphate buffered saline (PBS) followed by fixation with 4 % PFA at room temperature for 20 minutes. After fixation brain was carefully removed and sectioned with a vibratome at 40 μm thickness and all coronal midbrain sections were collected. Cresyl-violet staining was performed as described by Paul et al. (36). Briefly, samples were mounted on Menzel SuperFrost Ultra Plus™ slides (Thermo Fisher Scientific) and allowed to dry for 24 hours on room temperature. Slides were processed together and the same batch of reagents were used. Samples were demyelinated using increased ethanol containing solutions (2 min each) and Xylene (5 min), followed by the return to aqueous phase (30 sec each step). Cresyl-violet staining was performed using 0.1 % solution for 3 minutes. Clearing of the samples was carried out in 50 % ethanol for 15 minutes. Slides were fixed in xylene and mounted overnight with DPX Mounting medium (Sigma-Aldrich GmbH).

### Tyrosine hydroxylase (TH) immunohistochemistry

Mice were sacrificed by CO_2_ inhalation and the chest cavity was opened for perfusion with 4 °C phosphate buffered saline (PBS) followed by fixation with 4 % PFA at room temperature for 20 minutes. After fixation brain was carefully removed and sectioned with a vibratome at 40 μm thickness and all coronal midbrain sections were collected. Ten consecutive sections were used for immunostaining. Sections were processed together and the same batch of reagents were used. Endogenous peroxidase activity was blocked by 0.3 % H_2_O_2_ in methanol for 20 min. To reduce non-specific binding, Vector blocking solution (2.5 % normal horse serum) was applied for 2 h at room temperature. Sections were incubated with anti-tyrosine hydroxylase antibody (Sigma-Aldrich, Cat. No. AB152) overnight at 4 °C. Following, the secondary antibody (The ImmPRESS Universal Antibody Kit, anti-mouse/rabbit) and ImmPACT DAB (both purchased from Vector Laboratories, Burlingame, CA) was applied according to the manufacturer’s instructions. The sections were dried on glass slides and coverslipped with ProLong™ Gold Antifade Mountant (Thermo Fisher Scientific, Waltham, MA, USA).

### Immunofluorescent labeling and confocal laser imaging

Animals were sacrificed and brain sections have been prepared as described previously. Samples were washed three times with PBS and incubated in PBS containing 5 % BSA and 0.3 % Triton X-100 for 2 hours. Samples were randomly divided into groups and were incubated overnight at 4 °C in the same buffer containing anti-CD68 antibody (1:250), anti-Iba1 antibody (1:500) together with anti-phospho-p38 MAPK (T180/Y182) antibody (1:250). Samples from the same animal were incubated overnight at 4 °C in the same buffer containing anti-Iba1 antibody (1:500) together with anti-p38 MAPK antibody (1:250) to test whether treatment influences the basal expression of p38 MAPK. Next, a second set of experiments were performed to test P2Y_12_-receptor expression, where samples were incubated overnight at 4 °C in the same buffer containing anti-CD68 antibody (1:250) together with anti-P2Y_12_-receptor antibody (1:1000). To validate the anti-P2Y_12_-receptor antibody specificity, samples from wild-type control and P2Y_12_R-KO mice were harvested and prepared as described. Samples were incubated overnight at 4 °C in the PBS containing 5 % BSA and 0.3 % Triton X-100 buffer supplemented with anti-Iba1 antibody (1:500) together with anti-P2Y_12_-receptor antibody (1:1000). After overnight incubation, samples were washed in PBS and incubated with the appropriate AlexaFluor-564 conjugated antibody and AlexaFluor-488 conjugated secondary antibody (Molecular Probes; 1:200) as well as with Hoechst 33342 (Thermo Fisher Scientific; 1:10000) for 1 hour avoiding exposure to light. Samples were mounted in ProLong™ Gold Antifade Mountant (Thermo Fisher Scientific) overnight. Immunofluorescent signal was analyzed using a Nikon Eclipse Ti-E inverted microscope (Nikon Instruments Europe B.V., Amsterdam, The Netherlands), and a C2 laser confocal system.

### Post-mortem human brain tissue

Brains of 4 patients with Alzheimer’s disease and 2 patients with Parkinson’s disease dementia, were removed within 3 days after death and immersion fixed in 4 % paraformaldehyde. Brain dissection, macroscopic description, regional sampling, tissue processing and staining were done following standard protocols as described in earlier (66) including BrainNet Europe and Brains for Dementia Research UK. Briefly, dissected and paraffin embedded samples from the middle frontal gyrus (Brodmann area 9) and striatum, respectively, were selected for this study, and 7 μm thick sections were cut on a sledge microtome. Sections were routinely de-waxed, blocked for endogenic peroxidase activities in ethanol containing 3% (v/v) H_2_O_2_ for 30 minutes, and heat-treated in appropriate antigen retrieval buffer solutions using a household microwave oven (5 min at 800 W, 2 × 5 min at 250 W). Non-specific epitopes were blocked with 1% (v/v) bovine serum albumin dissolved in Tris-buffered saline (TBS pH=7.4) for one hour. The sections were incubated with the primary antibodies in 1:500 concentration overnight at 4 °C. Detection was processed using biotin-free secondary antibodies (MACH 4 Universal HRP-Polymer, Biocare Medical LLC, Cat. No. M4U534). For visualization, 3,3’-diaminobenzidine tetrahydrochloride (DAB) reagent (Biocare Medical LLC, Cat. No. DB801) was applied. Harris’ haematoxylin was used to perform nuclear counterstain. After dehydration the sections were covered with Sclencell medium and Mountex coverslip.

### Determination of [^3^H]dopamine release ([^3^H]DA)

Experiments were performed on young adult (2-3 months) male wild-type and P2Y_12_ receptor knockout (*P2ry12^−/−^*) mice. The [^3^H]DA release experiments were conducted using the method with slight modifications described in our previous papers (35). Briefly, the mice were anaesthetized under light CO_2_ inhalation, and subsequently decapitated. The striatum was dissected in ice-cold Krebs solution saturated with 95 % O_2_ and 5 % CO_2_, sectioned (400-μm-thick slices) using a McIlwain tissue chopper and incubated in 1 ml of modified Krebs solution (113 mM NaCl, 4.7 mM KCl, 2.5 mM CaCl_2_, 1.2 mM KH_2_PO_4_, 1.2 mM MgSO_4_, 25.0 mM NaHCO_3_, and 11.5 mM glucose), pH 7.4, in the presence of 5 μCi/ml [^3^H]dopamine, specific activity 60 Ci/mmol; ARC, Saint Louis, MO, USA) for 45 min. The medium was bubbled with 95 % O_2_ and 5 % CO_2_ and maintained at 37 °C. After loading, the slices were continuously superfused with 95 % O_2_ and 5 % CO_2_-saturated modified Krebs solution (flow rate: 0.7 ml/min). After a 90 min washout period to remove excess radioactivity, perfusate samples were collected over 3 min periods and assayed for tritium content. The temperature was strictly kept at 37 °C. At 6 min after the start of the collection, the slices were subjected to a 3 min perfusion of the Na^+^-channel activator veratridine (5 μM) and then changed to normal Krebs solution until the end of the collection period. In some experiments, two consecutive veratridine stimulus were applied 30 min apart (S1, S2) and the P2Y_12_R antagonist PSB 0739 (1 μM) was perfused 15 min before the second veratridine stimulation (S2). The effect of PSB 0739 on the veratridine-evoked [^3^H]DA release was expressed as the ratio of S2 over S1 and compared with control S2/S1 values. The radioactivity released from the preparations was measured using a Packard 1900 Tricarb liquid scintillation spectrometer, using Ultima Gold Scintillation cocktail. The release of tritium was expressed in Bq/g or as a percentage of the amount of radioactivity in the tissue at the sample collection time (fractional release). The tritium uptake in the tissue was determined as the sum of release + the tissue content after the experiment and expressed in Bq/g. For the evaluation of the basal tritium outflow the fractional release measured in the first 3 min sample under drug free conditions were taken into account. The veratridine-induced [^3^H]DA efflux calculated as the net release in response to the respective stimulus by subtracting the release before the stimulation from the values measured after stimulation (S1, FRS1).

### Animal models

Animals were housed under a 12-hour light-dark cycle under specific pathogen-free conditions, and had access to food and water *ad libitum*. All mice were backcrossed onto a C57BL/6N background at least 8 to 10 times, and experiments were performed with littermates as controls. Experiments were performed using wild-type (C57/Bl6N) and *P2ry12* gene-deficient (*P2ry12^−/−^*) male mice aged 8-14 weeks. Embryonic stem cells for *P2ry12^−/−^* knockout mice, B6;129-P2ry12^tm1Dgen^/H were purchased from Deltagen Inc. (San Matteo, CA, USA). Cloning and breeding strategy, and genotyping protocol has been described previously (44).

Two different experimental approach was used;

1. *in vivo acute MPTP model*. The acute induction of Parkinson’s disease was described in detail elsewhere (67); briefly, animals were randomly assigned into experimental groups. All animals received either intrathecal injection of PSB 0739 (0.3 mg / kg) or saline 18 hours prior to MPTP treatment. MPTP was injected (4 × 20 mg / kg intraperitoneally) 2 hours apart; behavioral tests were performed 2 hours and 24 hours after the last MPTP injection; while animals were sacrificed either 36 hours for cytokine levels assessment or 72 hours after the last MPTP treatment for measurement of biogenic amine content determined by HPLC-EC analysis.
2. *in vivo subchronic MPTP model*. Animals were treated daily dose of MPTP (20 mg / kg, intraperitoneally) or saline for five consecutive days; subsequently, animals were randomly divided into two groups, receiving daily intrathecal injections of PSB 0739 (0.3 mg / kg) or saline for four consecutive days (67). Alternatively, animals were treated with daily dose of MPTP (20 mg / kg, intraperitoneally) or saline for five consecutive days, followed by replacing the drinking water containing either fasudil (50 mg / kg body weight per day) or its vehicle for three weeks. Behavioral tests were performed before MPTP administration and 21 days later. Mice were sacrificed 21 days after last MPTP treatment and biogenic amine content were determined by HPLC-EC analysis.

### HPLC determination of nucleotides, catechol- and indoleamines content

Catechol- and indole amines, nucleotides (ATP, ADP, AMP) and adenosine, in extracts from brain tissue were determined using HPLC method. Mice were decapitated 3 or 21 days after the last MPTP or vehicle treatment, the striatum was dissected on ice and was snap frozen in liquid nitrogen. The tissue was ultrasonically homogenized with 200 μl of ice-cold 0.01 M perchloric acid solution containing theophylline (as an internal standard) at a concentration of 10 μM and 0.5 mM sodium metabisulfite (antioxidant of biogenic amines). The tissue extract was centrifuged at 3510 × g for 10 min at 4 °C and the pellet was saved for protein measurement according to Lowry et al. (68). Perchloric anion from the supernatant was precipitated by 1 M potassium hydroxide, the precipitate was then removed by centrifugation. The sample extracts were kept at −20 °C until analysis. Quantification of nucleotides and biogenic amines from tissue was performed by online column switching separation. ACE Ultra Core Super 5 μm particle size packed columns from A.C.T.L. (Scotland) were used for analysis. The phenylhexyl packed (7.5 cm × 2.1 mm ID) column was used for online solid phase extraction (SPE). Upon completion of sample enrichment and purification the separation was continued by connecting the analytical C-18 (150 × 2.1 mm) column. The flow rate of the mobile phases [“A” 10 mM potassium phosphate, 0.25 mM EDTA “B” with 0.45 mM octane sulphonyl acid sodium salt, 8 % acetonitrile (v/v), 2 % methanol (v/v), pH 5.2] was 350 or 450 μl / min, respectively in a step gradient application (69). The enrichment and stripping flow rate of buffer [10 mM potassium phosphate, pH 5.2] was during 4 min and the total runtime was 55 min. The HPLC system used was a Shimadzu LC-20 AD Analytical & Measuring Instruments System, with an Agilent 1100 Series Variable Wavelength Detector and BAS CC-4 amperometric detector in a cascade line. The detection of nucleotides and the internal standard (theophylline) was performed at 253 nm wavelengths by UV and biogenic amines were determined electrochemically at an oxidation potential of 0.73 V. Concentrations were calculated by a two-point calibration curve internal standard method: (Ai × f × B)/(C × Di × E) (Ai: Area of nucleotide or biogenic amine component; B: Sample volume; C: Injection volume; Di: Response factor of 1 pmol biogenic amine and 1 nmol nucleotide standard; E: Protein content of sample; f: factor of Internal Standard (IS area in calibration / IS area in actual)). The data was expressed as pmol / mg protein, unless stated otherwise. Statistical analysis was performed using the TIBC Statistical Program to assess normal distribution of all continuous variables and the nonparametric Kolmogorov-Smirnov test was used to test statistical differences. Where the measured variables met the normality assumption parametric tests were performed. The threshold for statistical significance was set at p<0.05.

### Behavioral analyses

Rotarod test was performed to assess motor coordination on the IITC (Woodland Hills, CA, USA) Rotarod Apparatus. The modified protocol used was described by Shiotsuki (70). Briefly, the tests are performed on an 8 cm diameter rotating rod 25 cm above the base of the apparatus rotating with fixed speed (10 rpm) in order to obtain a steep learning curve, consequently this protocol is superior in the assessment of motor skill learning rather than maximal gait performance. Motor coordination of animals was tested for 180 seconds. Acclimatization to the device was performed for 2 consecutive days before the start of the experiment. Regarding the acute MPTP treatment model, baseline latencies to fall were determined 1 hour before drug administration, followed by either PSB 0739 (0.3 mg / kg) treatment or its vehicle. The falling latency was measured at 6 and 24 hours after the final MPTP treatment. Regarding the subchronic MPTP model, baseline values were obtained 1 hour prior to the start of MPTP treatment or its vehicle on day 1. The latency time to fall was again measured 21 days after the last MPTP administration.

### Flow cytometry

Brain samples were collected 24 hours after the last intraperitoneal injection of MPTP (20 mg / kg) or saline. Mice were perfused transcardially with 0.1 M phosphate buffered saline (PBS), and substantia nigra and striatum were homogenized in RIPA lysis buffer (150 mM NaCl, 50 mM Tris-HCl (pH 7.4), 5 mM EDTA, 0.1 % (w/v) SDS, 0.5 % sodium deoxycholate and 1 % Triton X-100) supplemented with protease inhibitors (71, 72). After centrifugation (16000 × g for 20 min at 4 °C), supernatants were collected and protein concentration were measured using BCA Protein Assay Kit (Thermo Fisher Scientific, Pierce). Concentration of IL-1β, IL-6, IL-10 and TNFα were measured using BD Cytometric Bead Array Flex Sets (BD Biosciences). Measurements were performed on a BD FACSVerse flow cytometer, and data were analyzed using the FCAP Array version 5 software (Soft Flow). Cytokine concentrations of brain tissue were normalized to total protein levels measured. The cytokine levels are expressed as pg / mg total protein, unless stated otherwise.

### Statistics

Statistical analysis was performed using the GraphPad Prism software v.6.07 from GraphPad Software Inc. (La Jolla, CA, USA). Values are presented as mean ± SEM; *n* represents the number of independent experiments. Probability distribution of all continuous variables was performed; nonparametric data were analyzed using Kolmogorov-Smirnov test, whereas in case of normally distributed data, statistical analysis between two groups were performed with an unpaired two-tailed Student’s *t* test, while multiple group comparisons were analyzed with one-way ANOVA followed by Tukey’s post-hoc test, unless stated otherwise, and comparisons between multiple groups at different time points were performed using two-way ANOVA followed by Bonferroni’s post-hoc test. A *p* value of less than 0.05 was considered to be statistically significant.

### Study approval

All procedures involving animal care and use in this study were performed in accordance with the Institutional Ethical Codex and the Hungarian Act of Animal Care and Experimentation guidelines (40/2013, II.14), which are in accordance with the European Communities Council Directive of September 22, 2010 (2010/63/EU). The Animal Care and Experimentation Committee of the Institute of Experimental Medicine and the Animal Health and Food Control Station, Budapest, have also approved all experiments (PEI/001/776-6/2015). Experimental animals were treated humanely, all efforts were made to minimise animal suffering and reduce numbers of experimental animals. Animal studies are reported in compliance with the ARRIVE guidelines. The exact number and genotype of experimental animals used are indicated in their respective figures legend. Sample size was calculated as described previously (73), and was estimated based on a pilot study of MPTP and vehicle treated mice. Human brain tissue was obtained from patients who died from causes not linked to brain diseases, and did not have a history of neurological disorders (ethical approval ETT TUKEB 31443/2011/EKU [518/PI/11]), and from patients with Parkinson’s disease (ethical approval ETT-TUKEB 62031/2015/EKU, 34/2016 and 31443/2011/EKU [518/PI/11]). Informed consent was obtained for the use of brain tissue and for access to medical records for research purposes, and the use of tissue samples were in accordance with the Declaration of Helsinki.

## Author contributions

AI performed most of the in vitro and in vivo experiments, analyzed and discussed data, and wrote the manuscript. FG, LO, BV, AT, and LVM conducted experiments and acquired and analyzed data. MB performed HPLC analysis, acquired and analyzed data. TH, DB and ÁD discussed data. BS initiated and supervised the study, discussed data, and contributed to writing the manuscript. All authors read and commented on the manuscript.

## Acknowledgements

This work was supported by the Hungarian Research and Development Fund [grant number 116654, 131629]; Hungarian Brain Research Program [2017-1.2.1-NKP-2017-00002 to B.S.], the European Union’s Horizon 2020 Research and Innovation Programme under the Marie Skłodowska-Curie grant agreement No. 766124, and the Hungarian Academy of Sciences Premium Postdoctoral Research Program [PPD2019-20/2019-439. to A.I.]. The authors are grateful for Zsuzsanna Környei for providing BV-2 cell line, and Dóra Gali-Györkei and Mónika Borza for technical assistance.

---

**Supplemental Table 1.**
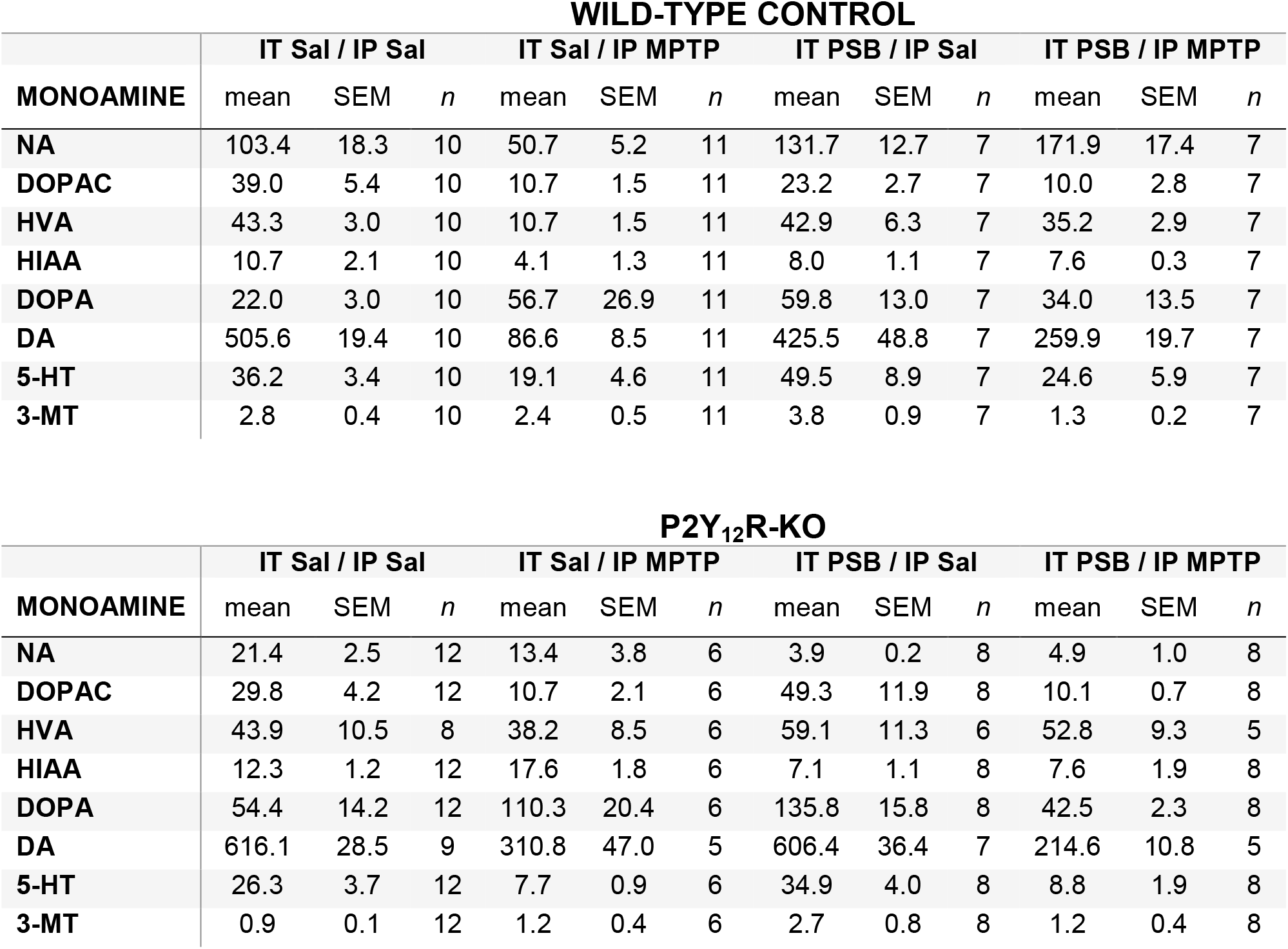
Monoamine concentration in WT and P2Y_12_R-KO mice. NA: Noradrenaline; DOPAC: 3,4-Dihydroxyphenylacetic acid; HVA: Homovanillic acid; HIAA: 5-Hydroxyindoleacetic acid; DOPA: 3,4-Dihydroxyphenylalanine; DA: Dopamine; 5-HT: 5-Hydroxytryptamine; 3-MT: 3-Methoxytyramine. Values of the mean and SEM are shown in pmol/mg protein.

**WILD-TYPE CONTROL**
| MONOAMINE | IT Sal / IP Sal |  |  | IT Sal / IP MPTP |  |  | IT PSB / IP Sal |  |  | IT PSB / IP MPTP |  |  |
| --- | --- | --- | --- | --- | --- | --- | --- | --- | --- | --- | --- | --- |
|  | mean | SEM | *n* | mean | SEM | *n* | mean | SEM | *n* | mean | SEM | *n* |
| NA | 103.4 | 18.3 | 10 | 50.7 | 5.2 | 11 | 131.7 | 12.7 | 7 | 171.9 | 17.4 | 7 |
| DOPAC | 39.0 | 5.4 | 10 | 10.7 | 1.5 | 11 | 23.2 | 2.7 | 7 | 10.0 | 2.8 | 7 |
| HVA | 43.3 | 3.0 | 10 | 10.7 | 1.5 | 11 | 42.9 | 6.3 | 7 | 35.2 | 2.9 | 7 |
| HIAA | 10.7 | 2.1 | 10 | 4.1 | 1.3 | 11 | 8.0 | 1.1 | 7 | 7.6 | 0.3 | 7 |
| DOPA | 22.0 | 3.0 | 10 | 56.7 | 26.9 | 11 | 59.8 | 13.0 | 7 | 34.0 | 13.5 | 7 |
| DA | 505.6 | 19.4 | 10 | 86.6 | 8.5 | 11 | 425.5 | 48.8 | 7 | 259.9 | 19.7 | 7 |
| 5-HT | 36.2 | 3.4 | 10 | 19.1 | 4.6 | 11 | 49.5 | 8.9 | 7 | 24.6 | 5.9 | 7 |
| 3-MT | 2.8 | 0.4 | 10 | 2.4 | 0.5 | 11 | 3.8 | 0.9 | 7 | 1.3 | 0.2 | 7 |

**Supplemental Table 1. Monoamine concentration in WT and P2Y<sub>12</sub>R-KO mice**
| MONOAMINE | IT Sal / IP Sal |  |  | IT Sal / IP MPTP |  |  | IT PSB / IP Sal |  |  | IT PSB / IP MPTP |  |  |
| --- | --- | --- | --- | --- | --- | --- | --- | --- | --- | --- | --- | --- |
|  | mean | SEM | *n* | mean | SEM | *n* | mean | SEM | *n* | mean | SEM | *n* |
| NA | 21.4 | 2.5 | 12 | 13.4 | 3.8 | 6 | 3.9 | 0.2 | 8 | 4.9 | 1.0 | 8 |
| DOPAC | 29.8 | 4.2 | 12 | 10.7 | 2.1 | 6 | 49.3 | 11.9 | 8 | 10.1 | 0.7 | 8 |
| HVA | 43.9 | 10.5 | 8 | 38.2 | 8.5 | 6 | 59.1 | 11.3 | 6 | 52.8 | 9.3 | 5 |
| HIAA | 12.3 | 1.2 | 12 | 17.6 | 1.8 | 6 | 7.1 | 1.1 | 8 | 7.6 | 1.9 | 8 |
| DOPA | 54.4 | 14.2 | 12 | 110.3 | 20.4 | 6 | 135.8 | 15.8 | 8 | 42.5 | 2.3 | 8 |
| DA | 616.1 | 28.5 | 9 | 310.8 | 47.0 | 5 | 606.4 | 36.4 | 7 | 214.6 | 10.8 | 5 |
| 5-HT | 26.3 | 3.7 | 12 | 7.7 | 0.9 | 6 | 34.9 | 4.0 | 8 | 8.8 | 1.9 | 8 |
| 3-MT | 0.9 | 0.1 | 12 | 1.2 | 0.4 | 6 | 2.7 | 0.8 | 8 | 1.2 | 0.4 | 8 |

**Supplemental Figure 1.**
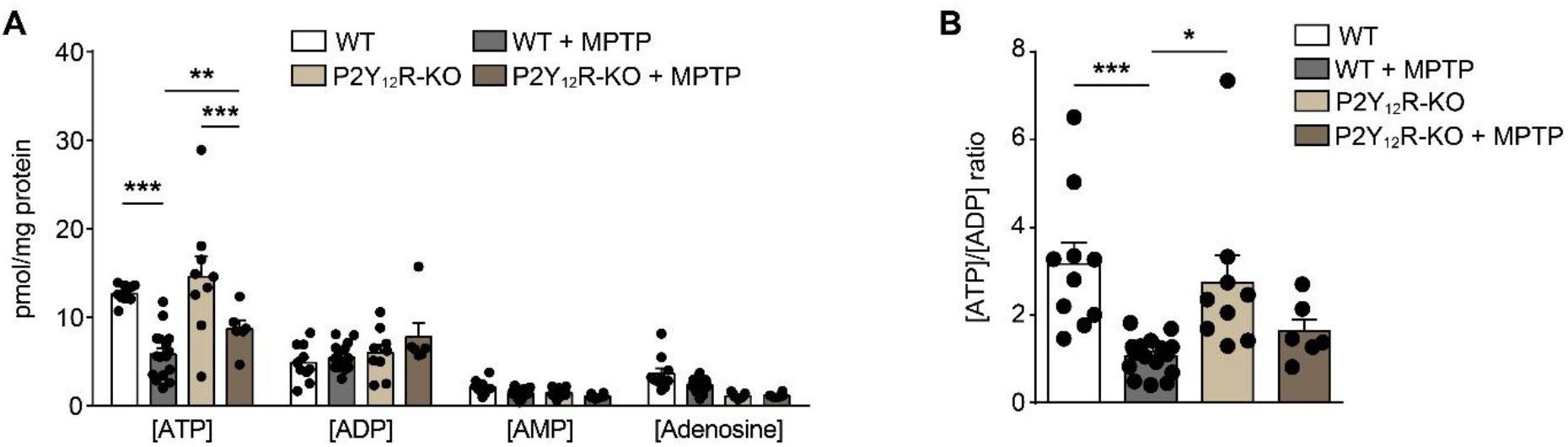
Nucleotide and adenosine levels measured in the striatum from wild-type and P2Y_12_R-KO mice after vehicle or MPTP-treatment. **(A)** WT or P2Y_12_-KO mice were treated with 4 × 20 mg / kg MPTP or its vehicle as indicated. Concentration of ATP, ADP, AMP and adenosine were determined from striatum samples 72 hours after last MPTP administration (n=6-16). **(B)** Ratio of ATP and ADP concentration (n=6-16). Data represent the mean ± SEM; *, *p* ≤ 0.05; **, *p* ≤ 0.01; ***, *p* ≤ 0.001 (two-way ANOVA, with Tukey’s *post-hoc* test (A, B)).

**Supplemental Figure 2.**
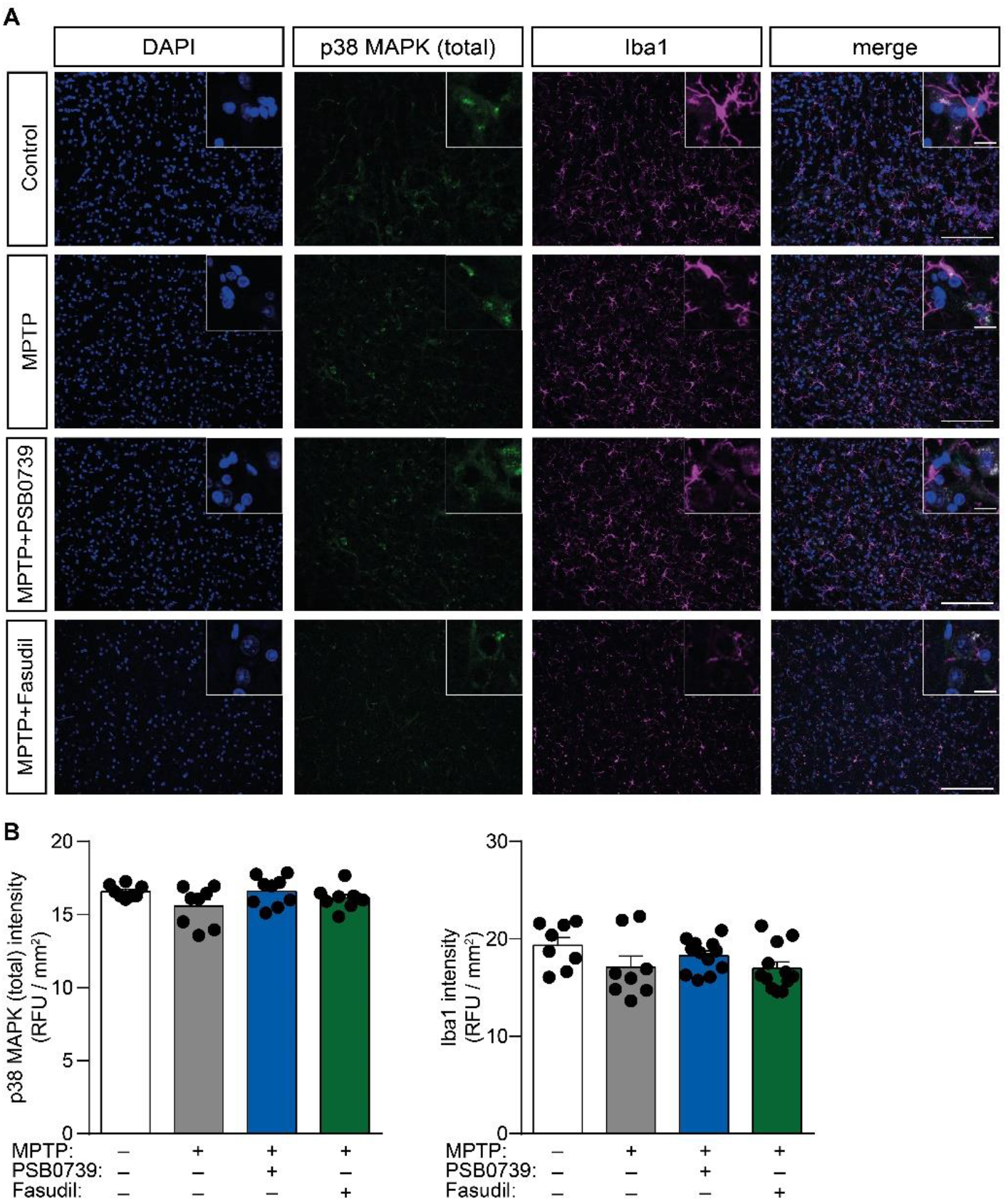
p38 MAPK total protein levels are unchanged in mice after MPTP-treatment. **(A-B)** WT mice were treated with 20 mg / kg MPTP daily for five consecutive days, followed by treatment with 0.3 mg / kg PSB 0739, 50 mg / kg body weight per day fasudil or its vehicle. Shown are representative immuno-confocal microscopy images of brain slices isolated from WT mice stained with antibodies directed against p38 MAPK (total; green), Iba1 (magenta), DAPI (blue) and overlay image (merge). Scale bar: 100 μm, corresponds to 20 μm inset **(A).** Quantification of p38 MAPK (total) fluorescence intensity (left panel) and Iba1 fluorescence intensity (right panel) (n=8-9, left panel; n= 8-12, right panel) **(B)**. Data represent the mean ± SEM; *, *p* ≤ 0.05; (two-way ANOVA, with Bonferroni’s *post-hoc* test (B)).

**Supplemental Figure 3.**
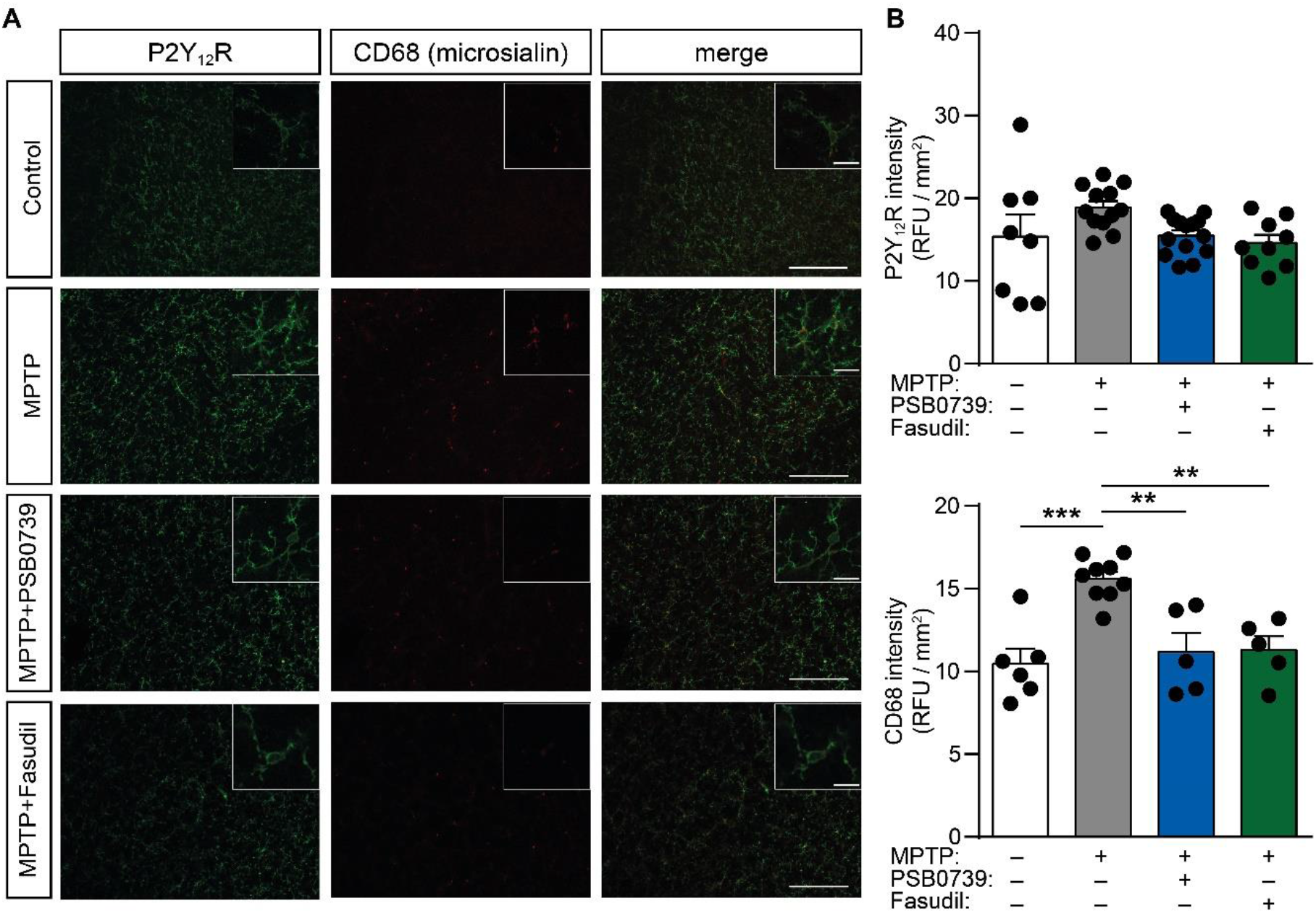
P2Y_12_R expression levels are unchanged in mice after MPTP-treatment. **(A-B)** WT mice were treated with 20 mg / kg MPTP daily for five consecutive days, followed by treatment with 0.3 mg / kg PSB 0739, 50 mg / kg body weight per day fasudil or its vehicle. Shown are representative immuno-confocal microscopy images of brain slices isolated from WT mice stained with antibodies directed against P2Y_12_R (green), CD68 (microsialin, red) and overlay image (merge). Scale bar: 100 μm, corresponds to 20 μm inset **(A)**. Quantification of P2Y_12_R fluorescence intensity (upper panel) and CD68 fluorescence intensity (lower panel) (n=8-16, upper panel; n= 5-9, lower panel) **(B)**. Data represent the mean ± SEM; **, *p* ≤ 0.01; ***, *p* ≤ 0.001 (two-way ANOVA, with Bonferroni’s *post-hoc* test (B)).

**Supplemental Figure 4.**
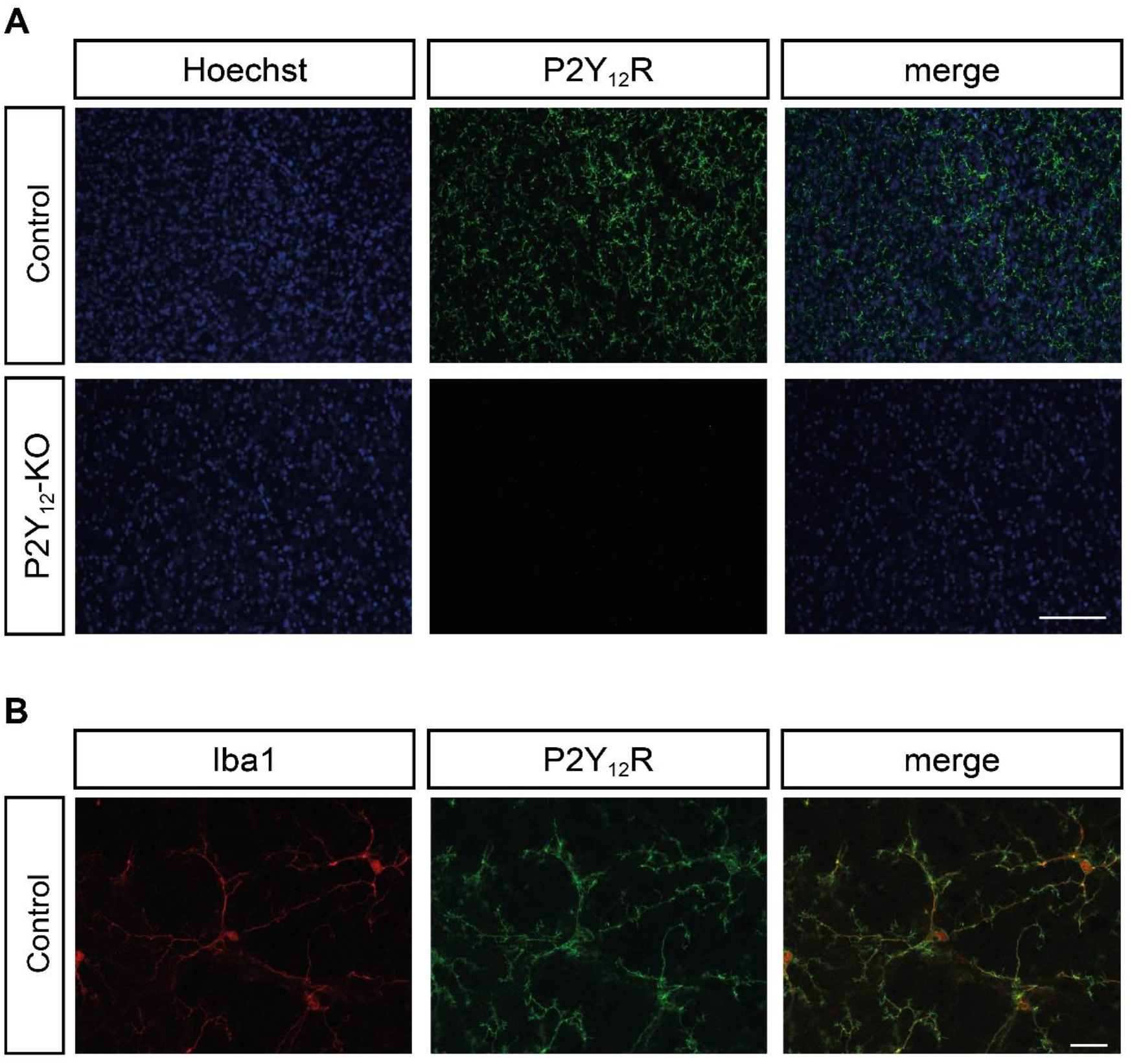
Validation of the P2Y_12_R antibody specificity. **(A)** Shown are representative immuno-confocal microscopy images of brain slices isolated from WT and P2Y_12_R-KO mice stained with antibodies directed against nuclei (Hoechst, blue), P2Y_12_R (green) and overlay image (merge). Scale bar: 100 μm. **(B)** Representative immuno-confocal microscopy images of brain slices isolated from WT mice stained with antibodies directed against Iba1 (red), P2Y_12_R (green) and overlay image (merge). Scale bar: 20 μm.

